# Inversion polymorphism in a complete human genome assembly

**DOI:** 10.1101/2022.10.06.511148

**Authors:** David Porubsky, William T. Harvey, Allison N. Rozanski, Jana Ebler, Wolfram Höps, Hufsah Ashraf, Patrick Hasenfeld, Human Pangenome Reference Consortium (HPRC), Human Genome Structural Variation Consortium (HGSVC), Benedict Paten, Ashley D. Sanders, Tobias Marschall, Jan O. Korbel, Evan E. Eichler

## Abstract

The completion of the human genome significantly improved our ability to discover and interpret genome copy number variation. In order to understand its impact on the characterization of inversion polymorphisms, we remapped data from 41 human genomes and 10 new samples against the telomere-to-telomere (T2T) reference genome as compared to the standard GRCh38 reference. Our analysis shows a ~21% increase in sensitivity identifying and improving mapping of 63 inversions. We further identify 26 misorientations within GRCh38, and show that the T2T reference is three times more likely to represent the correct orientation of the major human allele. As a result, we report a significant bias for inversions accumulating within the pericentromeric regions of specific chromosomes and show that functional annotations around inverted regions, such as topological-associated domains, can be better interpreted.

## Background

A gapless telomere-to-telomere (T2T) assembly of a human genome (T2T-CHM13) was recently released (Nurk et al. 2021). The complete reference newly resolved >240 Mbp of sequence not previously represented in GRCh38 improving the discovery of single-nucleotide variants (Aganezov et al. 2022) and copy number variants (Vollger et al. 2022). Compared to other classes of variation, the detection of balanced events such as inversions is particularly challenging (Jarvis et al. 2022). This is because most inversions are copy number neutral and are associated with repetitive DNA (Kidd et al. 2008; Porubsky et al. 2020; Porubsky, Höps, et al. 2022). This is especially true for the largest events that are most frequently flanked by long and highly identical segmental duplications (SDs), which themselves are copy number polymorphic among different individuals. Even among existing high-quality long-read genome assemblies, large inversion polymorphisms are often missed or incorrectly represented (Porubsky, Vollger, et al. 2022). While various approaches have been developed over the years to detect inversions, the Strand-seq platform remains among the most sensitive (Porubsky, Höps, et al. 2022; Hanlon, Lansdorp, and Guryev 2022). Accurate detection of inversions is critical for understanding human variation and disease because recurrent inversions have been shown to associate with regions prone to rearrange and cause neurodevelopmental disease (Osborne et al. 2001; Koolen et al. 2006; Cáceres et al. 2007; Zody et al. 2008; Cooper et al. 2011; Coe et al. 2014; Mohajeri et al. 2016).

The T2T-CHM13 assembly has been put forward as an improved human reference genome over the current incomplete GRCh38 and GRCh37 references. We sought to assess the potential advantage of detecting inversions on this new reference when compared to GRCh38 and whether it would significantly alter our understanding of the landscape and frequency of inversion polymorphism in the human genome. We specifically focused on the analysis of Strand-seq data generated previously from 41 human genomes (Porubsky, Höps, et al. 2022) and recalled inversions on the T2T-CHM13 reference. Our analysis uncovered orientation errors in the reference and identified novel inversion polymorphisms in previously inaccessible regions. We find that use of the new reference not only better represents minor and major inversion alleles but also provides new insights into enrichment within pericentromeric regions of the human genome. The work strongly suggests that this new reference should be adopted for human inversion discovery especially to discover new complex polymorphisms in more diverse human population samples—one of the goals of the Human Pangenome Reference Consortium (HPRC) (Wang et al. 2022).

## Results

### More accurate and complete inversion discovery with T2T reference

Previously, we generated Strand-seq data from 41 samples from the 1000 Genomes Project. Using the same algorithm applied to GRCh38, we remapped the Strand-seq data to T2T-CHM13 (v1.1) and combined it with both Bionano Genomics and assembly-based approaches to detect inversions (Porubsky, Höps, et al. 2022) (**Methods**). With this reanalysis we identified in total 373 inverted regions, including 296 balanced inversions, 56 inverted duplications, and 21 complex events across the autosomes and chromosome X (**Fig. S1A, Table S1, Methods**). For the remainder of this study, we focus exclusively on analysis of 296 balanced inversions on autosomes and chromosome X (and refer to these as inversions or inversion polymorphisms (**Fig. 1A**). While we report a comparable number of inverted bases per chromosome (**Fig. S1B**), the T2T callset increases the overall sensitivity for inversion detection by ~21% (63 likely novel inversions). Concomitantly, the total number of inverted bases (considering balanced inversions only) increases by ~10.5 Mbp (82.8 Mbp) compared to GRCh38 (72.3 Mbp). In addition, the GRCh38 reference harbors 26 misorientations—defined here as any region where all 41 samples are homozygous inverted compared to the reference. In stark contrast, no misorientations are defined in the T2T-CHM13 genome, confirming its value as an improved reference (**Fig. S1B**). Between the two references, inversion counts differ for most human chromosomes (n=19) with the majority showing a net increase on T2T-CHM13 (n=13) (**Fig. S2**). Consistent with earlier analyses (Porubsky, Höps, et al. 2022), Strand-seq detected the greatest proportion of inverted base pairs exclusively detecting 82 inversions with median size ~144 kbp and corresponding to ~80 Mbp of sequence (**Fig. S3**).

**Figure 1:**
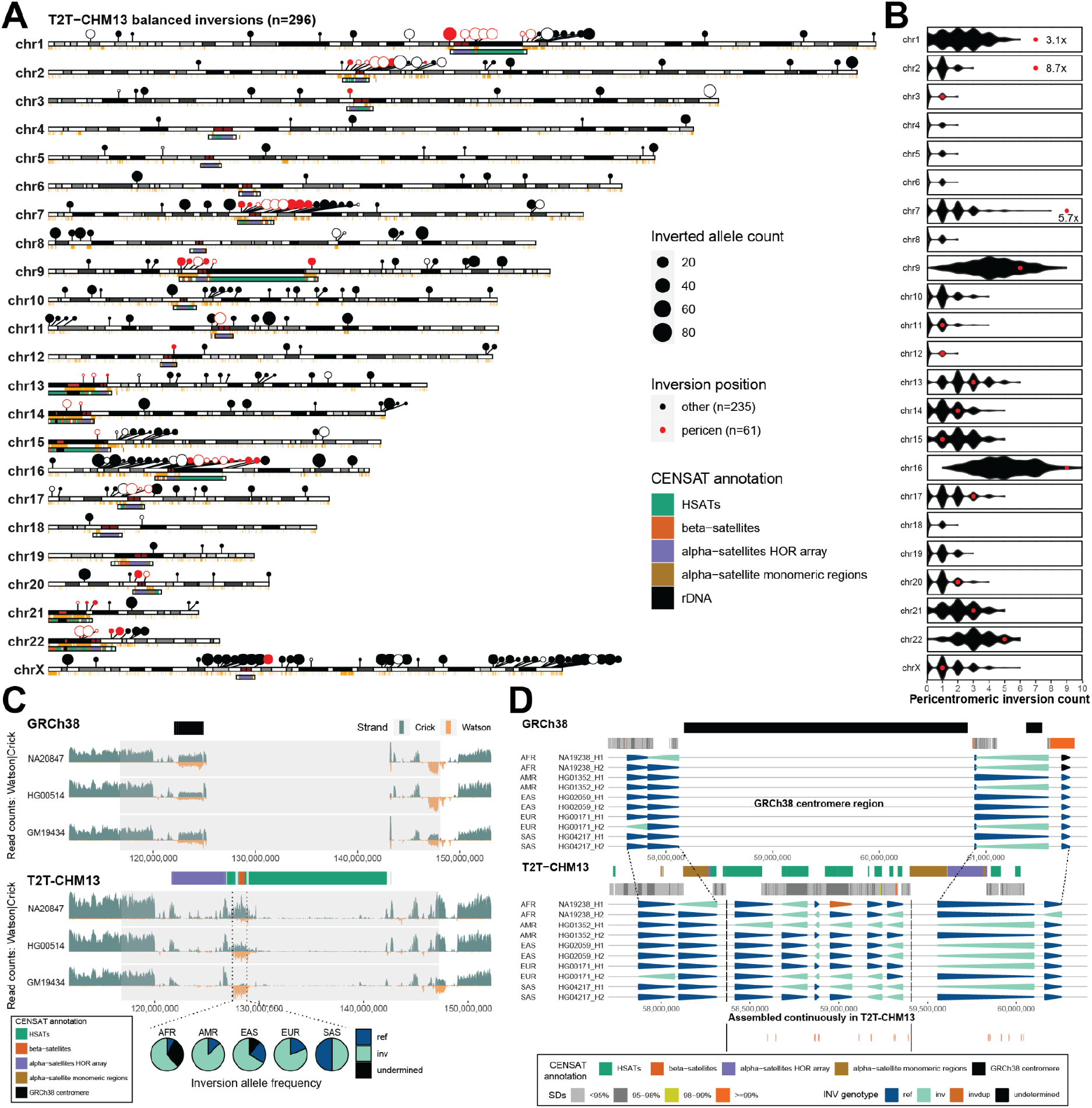
Inversion polymorphisms with respect to a complete T2T reference show pericentromeric bias. **A**) An ideogram showing the position and inverted allele frequency (dot size) of all balanced inversions from 41 human samples mapped to T2T-CHM13 reference (n=296). Inversions that fall within pericentromeric regions (CENSAT annotation, +/−1 Mbp) are shown as red dots (n=61) while other inversions are shown as black dots (n=235). Inversions with >=90% reciprocal overlap with nonsyntenic regions between GRCh38 and T2T-CHM13 or failed to map to the GRCh38 reference are highlighted as open circles (n=63). **B**) Permutation analysis shows pericentromeric enrichment for specific chromosomes. Permuted counts of pericentromeric inversions are shown as black violin plots as compared to observed counts (red dots). **C**) The read coverage profiles of Strand-seq data over a chromosome 1 centromeric region summarized as binned (bin size: 50 kbp step size: 10 kbp) read counts represented as bars above (teal; Crick read counts) and below (orange; Watson read counts) the midline with respect to centromere repeat annotation. Dotted lines highlight the novel centromeric inversion detected on chromosome 1 only with respect to T2T-CHM13. Note: equal coverage of Watson and Crick count represents a heterozygous inversion (one homologue inverted) while reads aligned only in the Watson orientation signify a homozygous inversion (both homologs inverted). Pie charts show frequency of inverted (bright blue) and directly oriented alleles (light blue) across all haplotypes (n=82) from all unrelated individuals (n=41) for a given centromeric inversion (dotted lines). **D**) A “backgammon” plot showing the inversion status of each defined region reported as colored arrowheads (dark blue - direct, bright blue - inverted, see the legend) for chromosome 7 region with respect to GRCh38 (chr7:57456486-61949954; top) and T2T-CHM13 (chr7:57700000-60400000; bottom). At the bottom there are novel genes (TEC) newly described in T2T-CHM13 reference.

### Novel inversions and pericentromeric enrichment

We identify 63 sites of putative novel inversions (**Fig. 1A**) when mapping to T2T-CHM13 (**Methods, Table S2**). Of these, 33 sites could be partially mapped to GRCh38 but share >=90% overlap with nonsyntenic regions present in the T2T-CHM13 but not GRCh38 reference and, thus, potentially represent structural differences between the references. In addition, there are 12 sites that failed to map to GRCh38, the majority of which are small (<1 kbp, n=8). Nevertheless, two of these unmapped sites, one on chromosome 7 and one on chromosome 17 are ~206 kbp and ~1.38 Mbp in size and have 39% and 52% overlap with nonsyntenic regions, respectively. Lastly, we define 18 sites that both failed to map to GRCh38 and have >=90% overlap with nonsyntenic regions and therefore are most likely novel (**Fig. S4**). Almost all of the nonsyntenic regions where the new inversions map correspond to pericentromeric sequence in T2T-CHM13—defined here as sequence +/−1 Mbp adjacent to rDNA or satellite DNA (**Methods**). Indeed, we find that 20.6% (61/296) of inversion polymorphisms are pericentromeric (**Fig. 1A**, **Methods**). The effect is particularly pronounced on chromosomes 1, 2 and 7 where we observe a three to eightfold enrichment (p < 0.05, Permutation Test with Bonferroni multiple testing correction) (**Fig. 1B, Fig. S5, Table S3, Methods**). Other chromosomes show more modest accumulation (i.e., chromosomes 9 and 16 with ~1.5-fold enrichments). We find 46% (28/61) of pericentromeric inversions associate with intrachromosomal SDs while another 26% (16/61) map to various classes of satellite DNA (**Fig. S6**). As an example, we identify a 1.3 Mbp inversion in the pericentromeric region of chromosome 1 that is completely absent from the GRCh38 reference (**Fig. 1C**). This large inversion represents the major allele in the human population (0.69 inverted allele frequency based on 82 analyzed haplotypes) (**Fig. 1C**). It is composed mostly of satellite repeats (human HSAT and beta satellites) and we predict inversion breakpoints fall within or nearby HSAT repeats (**Fig. S7**). We managed to confirm this inversion in four out of six HPRC assemblies of chromosome 1 centromere region (**Fig. S8**) and found notable variability in inversion size and its distance to proximal alpha satellite repeat array (**Fig. S9**). A second example includes a large 1 Mbp cluster of inversions mapping to the pericentromeric region of chromosome 7. In T2T-CHM13, this region is continuously assembled and contains six inverted loci that are either missing or likely misassembled in the GRCh38 reference (**Fig. 1D, Fig. S10**). The six satellite-associated inversions are polymorphic creating a diverse pattern of haplotypic structural diversity in the human population. Notably, there are multiple novel predicted genes (TECs) that remain to be validated and characterized (Nurk et al. 2021) (**Fig. 1D**).

### Improved annotation of inversion polymorphisms

As mentioned above, we identified 26 regions where GRCh38 differed in orientation with respect to T2T-CHM13, but all 82 human haplotypes supported the T2T-CHM13 configuration (**Fig. S11, Table S4**). While it is possible that these could be very low-frequency inversion polymorphisms, it is more likely that these simply represent misorientation errors. Many of these putative errors are large with median size of 16,306 bp (range: 488-2,346,462 bp). Excluding these likely misorientations, we find that T2T-CHM13 is much more likely to carry the major allele in the population. Specifically, we observe a threefold reduction of minor inversion alleles in T2T-CHM13 (n=11) compared to GRCh38 (n=33) (**Fig. S12**). Because these regions contain or map near protein-coding genes (**Fig. S12, Table S4**), these flips in orientation or changes in the major allele definition (**Fig. 2A**) can affect our interpretation of human genetic variation and functional annotation of the human genome. Such is the case for the melanoma antigen gene family cluster (*MAGE*) inversion polymorphisms mapping to the chromosome Xq28 region. In this region a minor (inverted) allele was originally reported in GRCh38 leading to the prediction of a series of nested inversions within a single haplotype (Porubsky, Höps, et al. 2022). However, with respect to T2T-CHM13, we can now report that the direct configuration represents the major allele (**Fig. S13**). Rather than nested inversions, we observe four independent inversion events utilizing distinct SD pairs and affecting different *MAGE* genes at various frequencies in the human population. Analysis of HPRC phased genome assemblies (**Fig. 2B, Methods**) confirms five distinct human structural configurations that result in inversion polymorphisms of different sizes and frequencies among human populations with H5 predicted to be ancestral based on structural similarity to chimpanzee (**Fig. S14**). Similarly, disease-associated regions such as the 16p12.1 microdeletion region associated with neurodevelopmental disabilities (Cooper et al. 2011; Coe et al. 2014; Bragin et al. 2014) are now properly configured. This region was reported to be misoriented in the GRCh38 reference (Antonacci et al. 2010) and is now correctly configured within the T2T-CHM13 reference (Nurk et al. 2021) (**Fig. S15**) helping to better distinguish long and short inversions in this region. Finally, because proximity ligation experiments such as HiC depend on correct order and orientation of the assembled sequence, the correction of these errors can affect functional genome annotation. Such is the case when detecting topologically associated domains (TADs) at 16p12.1 that carry the long (GM20847) and short (HG02011) versions of an inversion at this locus (**Fig. 2C, Methods**). Here, the GRCh38 reference reports hard-to-interpret regional associations while T2T-CHM13 provides a much clearer picture of TADs that are in line with a presence of reported inversions (**Fig. 2C**).

**Figure 2:**
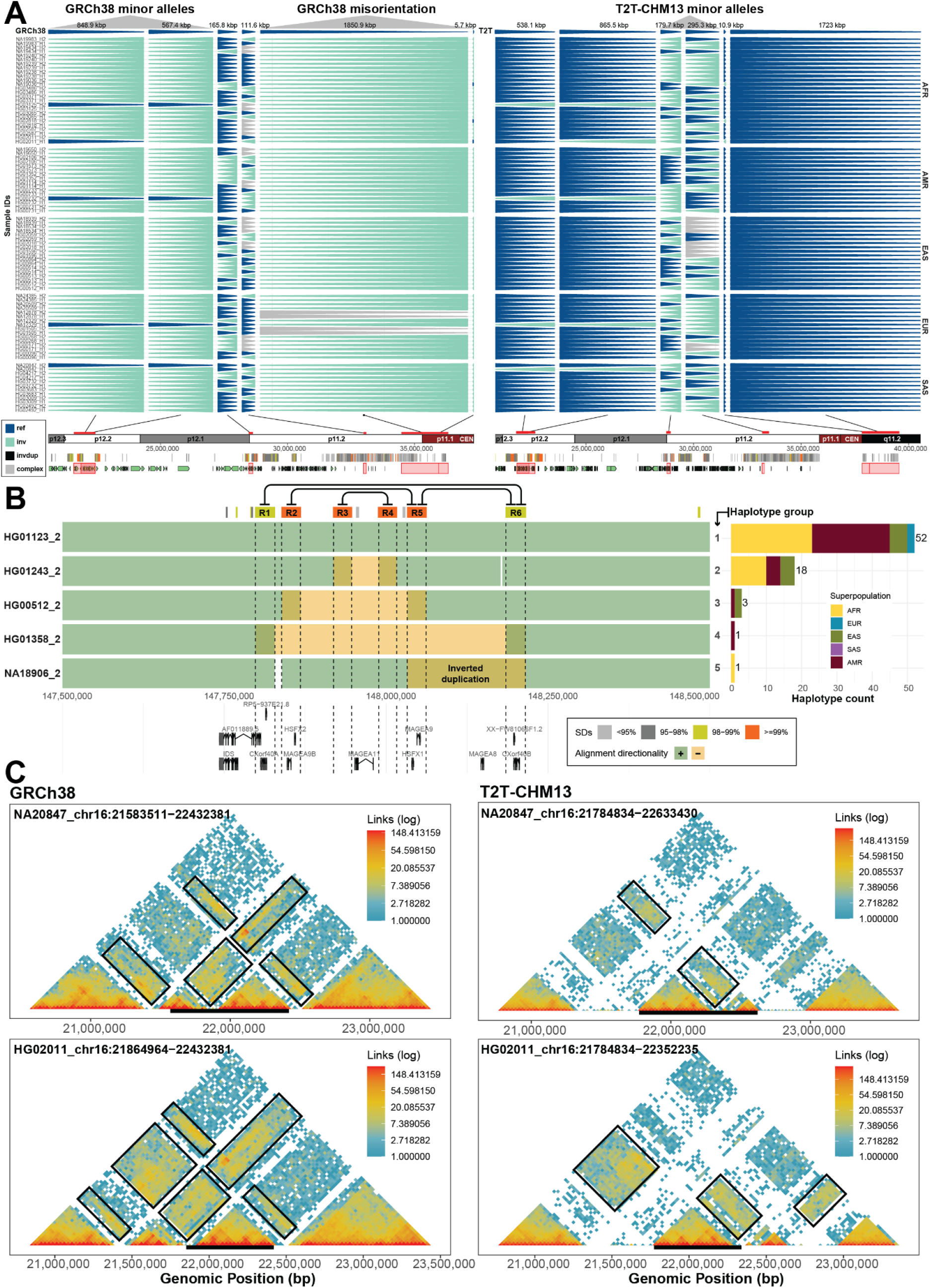
Improved representation of inversion polymorphism in T2T-CHM13 and interpretation of TADs. **A**) A “backgammon” plot for a 20 Mbp region across chromosome 16 depicting changes in the representation of major alleles as inverted (light blue) and direct orientation (dark blue) based on phased inversion genotypes reported with respect to GRCh38 and T2T-CHM13 reference genomes. In most cases, GRCh38 was either erroneous or represented the minor allele. **B**) Overlapping inversions on chromosome Xq28. Each row represents a unique human haplotype (haplotypes 1-5) of the Xq28 region visualized as a single human assembly aligned to T2T-CHM13 with directly orientated (green) and inverted (orange) segments displayed with respect to flanking segmental duplications (R1-6) that likely mediate the inversions (connecting lines) and underlying protein-coding genes. We use transparency to convey positions of overlapping alignments such as highlighted inverted duplication in haplotype 5. Barplot (right) shows the total counts of human haplotypes per haplotype group stratified by superpopulation. **C**) Contact matrices deduced based on Hi-C data mapped to GRCh38 and T2T-CHM13 reference are shown on the left and right side, respectively. We present contact matrices constructed for two samples (HG02011 and GM20847) with short and long versions of the inversion over the selected chromosome 16 region (black bar at the bottom). Intensity of contacts between proximal regions of the genome is represented by a heatmap colors from low level of contacts (blue) to high level of contacts (red). Regions with different levels of contact between two matrices are highlighted by black rectangles.

### Discovery of novel rare inversion polymorphisms and disease-associated rearrangements

As part of a quality control assessment during the development of the first phase of the HPRC (Porubsky, Vollger, et al. 2022; Liao et al. 2022), 10 additional Strand-seq datasets (**Data availability**) were generated from unrelated individuals for the HPRC. We applied these data to the T2T-CHM13 reference in an effort to discover additional rarer inversion polymorphisms. While the vast majority of inverted polymorphisms had been identified previously among the original 41 samples, we did identify five additional inversions (**Table S5**), including a novel structural configuration for the Xq28 *MAGE* gene cluster described above. The other novel inversions include large >1 Mbp events corresponding to chromosomes 15q25.2, 16p11.2-12.2, 16q22.1-23.1 and 22q11.21 (**Fig. 3A**). Interestingly, all but one of these rare inversion polymorphisms overlap a pathogenic copy number variant in the human population, strengthening our recent observation of disease association (Porubsky, Höps, et al. 2022). This includes a large inversion polymorphism encompassing one of the most common rearrangements associated with autism at chromosome 16p11.2, which maps to the DiGeorge/VCF syndrome critical region interval that has been extensively studied (Gebhardt et al. 2003; Vergés et al. 2017) and a multi-Mbp inversion corresponding to the Cooper syndrome region on chromosome 15q25.2. Using the HPRC assemblies (**Data availability**), we define eight structurally diverse haplotypes with various frequencies in human populations (**Fig. 3B**). We predict haplogroups 1 and 8 to be protected while haplogroups 4-7 are likely at increased risk of microdeletion/duplication of 15q25.2 region due to higher number of SD bases in direct orientation (**Fig. 3B, Fig. S16**). We successfully characterized the SD-associated inversion breakpoints of the inverted haplotype corresponding to haplogroup 6 with inversion breakpoint falling within the ~5 kbp region of nearly perfect homology (99.8% identical) (**Fig. S17**). Lastly, we report a massive ~4 Mbp inversion located at chromosome 16q22.1-23.1, a region previously linked to prostate cancer (Osman et al. 1997) where, to our knowledge, inversion has yet to be identified. Comparison to nonhuman primate data (Porubsky et al. 2020) and previous studies (Maggiolini et al. 2019), suggest that for three out of five events the rare, inverted configuration in the human population represents the ancestral orientation (**Fig. 3A**).

**Figure 3:**
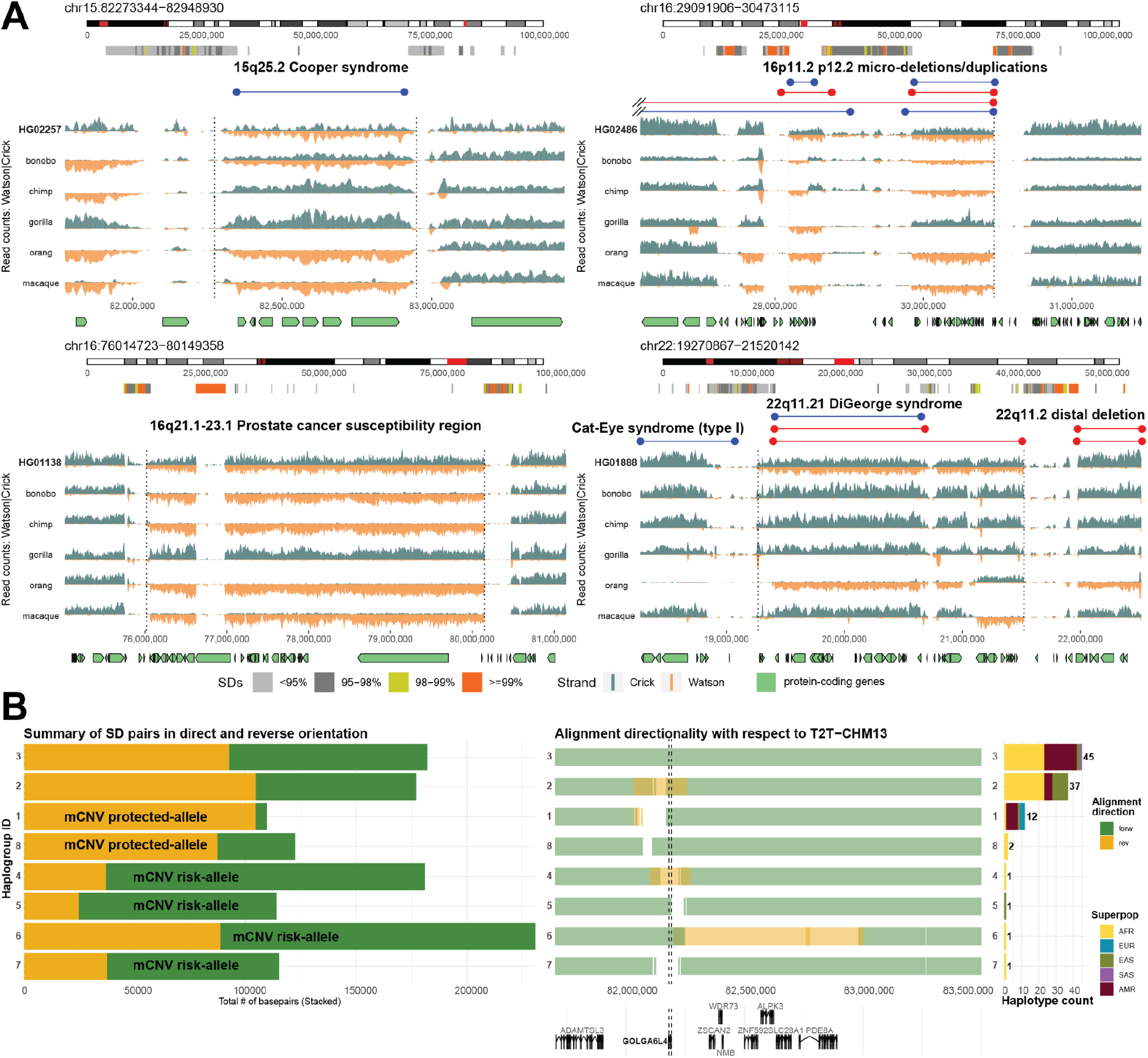
Rare inversion polymorphisms and disease-associated regions. **A)** Four disease-associated regions mapping to chromosomes 15q25.2, 16p11.2-12.2, 16q22.1-22.2, and 22q11.21 are depicted within chromosome-specific ideograms (red rectangle) with a zoom into the region flanked by segmental duplications (colored horizontal bars) and pathogenic duplication and deletion breakpoints highlighted in blue and red horizontal lines, respectively. Strand-seq data highlights rare heterozygous inversions (see **Fig. 1C** for detailed description) discovered in a human sample with respect to the status in different nonhuman primate species. Homozygous inversions are orange while homozygous teal represent homozygous direct orientations. **B**) Left plot summarizes the total number of base pairs for direct and reverse orientated SD pairs for each haplogroup (in rows) marked as likely protected or at risk for morbid copy number variation (mCNV) formation. Middle plot shows unique human haplotypes (haplotypes 1-8) of the 15q25.2 region visualized as a single human assembly aligned to T2T-CHM13 with directly orientated (green) and inverted (orange) segments. Underlying protein-coding genes from this region are shown below. Barplot (right) shows the total counts of human haplotypes per haplotype group stratified by superpopulation.

## Discussion

Previous studies have highlighted an increase in both specificity and sensitivity for single-nucleotide variant and copy number variant detection when using the T2T reference in lieu of GRCh38 (Aganezov et al. 2022; Vollger et al. 2022). Our results suggest the effect is the most pronounced for the discovery and characterization of inversion polymorphisms. The 20% gain in discovery stems in large part from the fact that inversions most strongly associate with repetitive DNA (Kidd, Graves, et al. 2010; Chaisson et al. 2019; Puig et al. 2020) and the addition of these previously inaccessible regions allows for their discovery by the mapping of Strand-seq data to these regions for the first time. In addition to these new discoveries, we provide further evidence that the T2T-CHM13 reference better represents the orientation of the major allele and identify 26 relatively large misorientations (total of 6.4 Mbp of sequence) in the original GRCh38 reference genome that have persisted for many earlier iterations of the human reference genome (**Fig. S18**). These results, thus, significantly improve our understanding of the landscape of inversion polymorphism in the human genome and argue that the T2T-CHM13 should be the preferential reference for future investigations into such human genetic variation.

Our analysis also revealed a greater propensity of polymorphic inversions to cluster within pericentromeric regions. However, this may not be surprising given that pericentromeric regions have been known for more than two decades to be hotspots for the accumulation of high-identity SDs (Eichler et al. 1995; She et al. 2004; Kidd, Sampas, et al. 2010; Altemose et al. 2022). Intrachromosomal SDs, in particular, drive the formation of many of the largest inversions via non-allelic homologous recombination (NAHR) and indeed nearly 50% of the pericentromeric inversions in this study have intrachromosomal SD pairs delineating their breakpoints (**Fig. S6**). Interestingly, we also observe a relatively high proportion (16%) of satellite-associated inversion polymorphisms, especially within selected pericentromeric regions where they appear to cluster creating considerable haplotypic diversity (**Fig. 1C,D**). Polymorphic inversions have the capacity to reduce recombination (Sturtevant 1917) and one possibility for reduced recombination across centromeres may be that the enrichment of such pericentromeric inversions at their flanks interferes with synapsis during meiosis. Alternatively, the reduced recombination may predate these structural features and instead promote the accumulation of large repeats promoting unequal crossover and inversion formation. As more genomes are characterized and these regions completely sequenced, it will be interesting to map recombination events and reconstruct haplotypes across these regions in relation to the massive structural differences.

Finally, our investigation into 10 more human genomes from the HPRC (Porubsky, Vollger, et al. 2022) continues to highlight the value of continued inversion polymorphism discovery. As previously reported (Porubsky, Höps, et al. 2022), the number of new inversions discovered is predictably low when compared to other forms of human variation due to an excess of common variation for this class of variant in the human population. Nevertheless, the additional rare polymorphisms we identified are >1 Mbp in length, spanning large swaths of genes and overlapping regions of genomic instability related to human disease. In particular, we recently demonstrated a fivefold association of recurrent inversion polymorphisms with recurrent genomic disorders among children with neurodevelopmental disorders (Porubsky, Höps, et al. 2022). One hypothesis is that the recurrent inversions are reshaping the architecture of the flanking SDs creating both protective and predisposed haplotypes to rearrangement as has been shown for a few loci (Zody et al. 2008; Kong et al. 2010; Steinberg et al. 2012). To address this, it will be critical to survey many more human genomes and to sequence resolve the large complex SDs flanking the inversion polymorphisms. Currently, methods such as trio-hifiasm fail to fully sequence resolve the many complex flanking SD regions or, in some cases, do not even identify the large inversion polymorphisms based on sequence assembly only. New methods, however, such as Verkko (Rautiainen et al. 2022) that couple both HiFi (high-fidelity PacBio) and ultra-long ONT (Oxford Nanopore) data show considerable promise in resolving a greater fraction of these regions. Once a large number of such haplotypes are fully sequenced and assembled, it will be possible to directly test whether predisposing and protective structural haplotypes exist by mapping sequencing data from patients with rearrangements to these new reference genomes (Itsara et al. 2009).

## Methods

### Strand-seq data generation and data processing

Strand-seq data were generated as follows. EBV-transformed lymphoblastoid cell lines from the 1KG (Coriell Institute) were cultured with BrdU (100 uM final concentration; Sigma, B9285) for 18 or 24 hours, and single isolated nuclei (0.1% NP-40 lysis buffer (Sanders et al. 2017)) were sorted into 96-well plates using the BD FACSMelody cell sorter. In each sorted plate, 94 single cells plus one 100-cell positive control and one 0-cell negative control were deposited. Strand-specific single-cell DNA sequencing libraries were generated using the previously described Strand-seq protocol (Falconer et al. 2012; Sanders et al. 2017) and automated on the Beckman Coulter Biomek FXp liquid handling robotic system (Sanders et al. 2020). Following 15 rounds of PCR amplification, 288 individually barcoded libraries (amounting to three 96-well plates) were pooled for sequencing on the Illumina NextSeq 500 platform (MID-mode, 75 bp paired-end protocol). The demultiplexed FASTQ files were aligned to the T2T-CHM13 reference assembly (v1.0 and v1.1) using BWA (version 0.7.15-0.7.17) for standard library selection. Aligned reads were sorted by genomic position using SAMtools (version 1.10) and duplicate reads were marked using sambamba (version 1.0). Low-quality libraries were excluded from future analyses if they showed low read counts (<50 reads per Mbp), uneven coverage, or an excess of ‘background reads’ (reads mapped in opposing orientation for chromosomes expected to inherit only Crick or Watson strands) yielding noisy single-cell data, as previously described (Sanders et al. 2017).

### Generation of inversion callset with respect to the T2T-CHM13 reference (v1.0)

In this study we applied the same multi-platform-based inversion discovery procedure as reported recently (Porubsky, Höps, et al. 2022). This procedure involves independent inversion discovery using PAV (long-read-based phased assemblies), Strand-seq (strand-specific short-read sequencing), and Bionano Genomics (optical mapping). We note that the inversion callset based on Strand-seq and Bionano Genomics underwent extensive manual curation in order to ensure high accuracy of a final inversion callset. Subsequently, independent inversion callsets were merged into a nonredundant inversion callset using SV-pop (Audano et al. 2019; Ebert et al. 2021).

### Lifting inversion callset to the latest version of T2T-CHM13 reference (v1.1)

Since the original inversion callset was done with respect to T2T-CHM13 v1.0, we decided to lift coordinates to v1.1 using liftOver. The only differences between v1.0 and v1.1 includes addition of missing rDNA and improved polishing within telomeres. To report inversion coordinates with respect to the latest version of T2T-CHM13 (v1.1, only difference with v2.0 is an addition of chromosome Y) reference, we used a command line version of UCSC liftOver tool (liftOver {input.bed} {input.chain} {output.bed} {output.unmapped}). We used publicly available liftOver chains ‘v1.0_to_v1.1.chain’ at https://s3-us-west-2.amazonaws.com/human-pangenomics/index.html?prefix=T2T/CHM13/assemblies/changes/v1.0_to_v1.1/. We attempted to lift all 374 detected sites. Of those, only one site genotyped as an inverted duplication (‘chr14-2842055-INV-181339’) positioned in chr14 rDNA failed to lift. Importantly, all sites (n=296) genotyped as balanced inversions were successfully lifted to T2T-CHM13 (v1.1/v2.0) coordinates. These coordinates will be used for all analyses reported in this paper (**Table S1**). Similarly, we used liftOver chains to translate coordinates from T2T-CHM13 to GRCh38, as was done for a complex region on chromosome 7 reported in Figure 1D; liftOver chain from T2T-CHM13 v2.0 to GRCh38 was downloaded from https://s3-us-west-2.amazonaws.com/human-pangenomics/index.html?prefix=T2T/CHM13/assemblies/chain/v1_nflo/chm13v2-grch38.chain.

### Mapping inversion coordinates to GRCh38

To translate coordinates of GRCh38 inversions into the T2T-CHM13 coordinate space, we decided to extract FASTA sequence from each inverted region and try to map such FASTA sequence onto the T2T-CHM13 reference using minimap2. This is because breakpoints of many inversions lie within SDs, which makes simple lifting of coordinates using liftOver difficult and results in many inverted regions to fail to lift. We mapped FASTA sequence extracted from inverted regions in GRCh38 to the T2T-CHM13 reference using minimap2 (version 2.24) with following parameters: -secondary=no --eqx -ax asm20. We filtered out alignments with mapping quality zero and alignments mapped to a different chromosome than the one FASTA sequence was extracted from. Inverted regions divided in multiple mappings were collapsed together, such that distance between subsequent mapping were no longer than 100 kbp. Lastly, we excluded mapped and collapsed ranges whose size was more than 50% larger or smaller than the original inversion range. Using this procedure, we were able to map 266 of all 296 balanced inversions in T2T-CHM13 callset. The same procedure was used when mapping inversion coordinates from GRCh38 to T2T-CHM13.

### Definition of likely novel inversions in T2T-CHM13

To define likely novel inversions detected with respect to T2T-CHM13, we set to investigate mappings of T2T-CHM13 inverted regions onto the GRCh38 reference (see section above) as well as previously defined nonsyntenic regions between T2T-CHM13 and GRCh38. The annotation of nonsyntenic regions in T2T-CHM13 with respect to GRCh38 was taken from the previous study (Vollger et al. 2022). We calculated the percent overlap between T2T-CHM13 inversions (n=296) and the list of nonsyntenic regions. Inversions with >=90% with the nonsyntenic regions were deemed as ‘nonsyntenic’ because their structure and relative orientation between T2T-CHM13 and GRCh38 might differ (**Fig. S4**). We marked inverted sites reported as nonsyntenic that also failed to map onto the GRCh38 reference as likely novel inversions in T2T-CHM13.

### Analysis of pericentromeric inversions

To define if an inversion lies within a peri-centromeric regions of the T2T-CHM13 assembly, we took a recently released annotation of centromeric repeats from https://s3-us-west-2.amazonaws.com/human-pangenomics/index.html?prefix=T2T/CHM13/assemblies/annotation/chm13.draft_v1.1.cenAnnotation.bed. Pericentromeric regions were defined as regions that include human satellites (‘hsat’ regions), alpha satellites (‘hor’ arrays), and rDNA with 1 Mbp of extra sequence at its flanks. Inversions that overlap (at least one base pair) with defined pericentromeric regions are considered as ‘pericentromeric’. Next, we used the R package regioneR (Gel et al. 2016) with its function ‘permTEST’ to perform permutation testing (n = 1,000 permutations) of pericentromeric inversions per chromosome. At each permutation, we randomized the position of each inversion per chromosome using regioneR’s function ‘randomizeRegions’ such that each inversion is assigned a random position along a given chromosome at each permutation. At each permutation, we counted the number of inversions overlapping with the pericentromeric region of any given chromosome. Due to multiple testing, we adjusted resulting p-values using Bonferroni correction. Subsequently, we evaluated the sequence composition of each inversion from pericentromeric regions (n=61). To do this, we calculated overlap of inverted bases with a set of genomic features, such as human satellites (HSATs), beta satellites, alpha satellites, monomeric regions, rDNA, and SD pairs. SD pairs were defined as intrachromosomal SDs that are no further apart than 5 Mbp. Inverted bases that do not overlap any of the above listed features were marked as ‘other’.

### Extraction of FASTA sequence from a region of interest

To extract FASTA sequence from a region of interest in T2T-CHM13 coordinates, we aligned available human assemblies from HPRC and HGSVC datasets to the T2T-CHM13 (v1.1) reference using minimap2 (version 2.24) with the following parameters: ‘-x asm20 --secondary=no -s 25000’. Next, we used rustybam (version 0.1.27) and its functionality called ‘liftover’ in order to subset alignments in PAF format to a region of interest. Then we used this subsetted PAF file in order to extract FASTA sequence using R package SaaRclust (Porubsky et al. 2021) and its function ‘regions2FASTA’. We extracted FASTA files only from assemblies that span the region of interest in a single continuous contig.

### Minor allele detection and misorientation validation

First, we mapped balanced inversions and putative misorientations in GRCh38 coordinates (n=330) to T2T-CHM13 using the procedure described above. Of the total 330 regions, 311 (281 balanced inversions and 30 misorientations) were successfully mapped to T2T-CHM13 coordinates. Next, we calculated the fraction of Watson (minus, negative strand) and Crick (plus, positive strand) reads mapped to each region in GRCh38 and T2T-CHM13 coordinates across all unrelated samples (n=41) used in this study. We required that each evaluated site include at least 20 mapped Strand-seq reads in both reference coordinates. Subsequently, a minor allele was defined as a region where there is at least 25% difference between Watson and Crick reads fraction mapped to GRCh38 and T2T-CHM13 for any given region. Also, we required that the ratio of Watson and Crick reads with respect to both references is no more than 25% different. The minor allele in GRCh38 is reported if the fraction of Crick reads (plus reads) is smaller than the fraction of Crick reads in T2T-CHM13 over the same region. This means that the majority of reads map in minus orientation across all unrelated samples while the majority of reads with respect to T2T-CHM13 map in direct (plus) orientation. Minor alleles in T2T-CHM13 were defined in an opposite manner as sites with the majority of reads mapped in minus orientation while for the same region GRCh38 counts mostly plus reads.

### Hi-C data analysis and visualization

To visualize Hi-C data, we first aligned short paired-end reads to the reference genome of interest (T2T-CHM13, v1.1). For this we used BWA (version 0.7.17) (Li and Durbin 2010) as follows: ‘bwa mem −5SP {input.ref} {input.pair1} {input.pair2}’. After the alignment we mark duplicate reads using ‘sambamba markdup’ (Tarasov et al. 2015) and sorted by query name as is standard for Hi-C analysis pipelines. Such aligned BAM files were processed using R package diffHic (Lun and Smyth 2015). First, we used the diffHic function ‘preparePairs’ in order to read in Hi-C alignments. At this step we filtered out reads with mapping quality less than 10 and any duplicate reads. Next, we used the diffHic function called ‘squareCounts’ in order to count the number of Hi-C interactions between two genomic bins of user-defined size. Lastly, the level of genomic interactions was visualized as diagonal squares colored by continuous heatmap colors on log^10^ scale.

### Detecting novel inversions using Strand-seq

In this study we added 10 additional samples (**Data availability**) where we called inversions using Strand-seq only (Sanders et al. 2016; Chaisson et al. 2019). A novel inversion was defined as inverted site not detected among 41 samples used to generate main inversion callset with respect to T2T-CHM13. Each newly detected inversion was checked for support using Strand-seq data from nonhuman primates to evaluate ancestral state of a given locus. Each newly detected inversion shows change in orientation in at least one nonhuman primate.

### Genome structural diversity of Xq28 and 15q25.2 regions

To analyze each region of interest (ROI) in more detail, we aligned full genome assemblies (**Data availability**) to the T2T-CHM13 reference. We next used rustybam (version 0.1.27) (10.5281/zenodo.6342176) and its ‘liftover’ functionality in order to subset assembly-to-reference alignments to a certain ROI. We select only assembled contigs with a complete span of the ROI such that contig boundaries are no further than 100 kbp from left and right ROI coordinates. We reverse complement assembly FASTA sequence in case the first and last contig alignment of at least 50 kbp is in minus orientation with respect to the reference. This is done to synchronize orientation among all FASTA files. We use such FASTA files to visualize alignment directionality with respect to the reference (T2T-CHM13 v1.1). In the case of the 15q25.2 region, we also align each FASTA file to itself using minimap2 (version 2.24) with the following parameters: ‘−x asm10 -c --eqx -D -P --dual=yes -r10,50’. We record a relative orientation (reverse or direct) of such self-alignments within each haplotype and calculate fraction and the total length of these alignments. This information is then used as a proxy to predict if an intervening region (between flanking self-alignments) is predisposed to inversion or CNV.

### Inversion breakpoint mapping

To map inversion breakpoints of a defined inversion at the 15q25.2 region, we selected the FASTA sequence of an inverted haplotype (HG02257_1) and direct haplotype from T2T-CHM13 corresponding to region chr15:81700000-83500000. Next, we aligned both inverted and direct haplotypes to themselves using minimap2 (version 2.24) in order to define pairs of identical sequences (SDs) within each haplotype. We selected only those pairs that were at least 500 kbp distance in order to obtain only those pairs that flank the inverted region (~675 kbp in size). We further selected those pairs that are in an inverted orientation with respect to each other. Lastly, we extracted FASTA sequence from such SD pairs for both inverted and direct haplotypes and continued with inversion breakpoint mapping as described in Porubsky et al. (2022).

## Acknowledgements

We thank Tonia Brown for assistance in editing this manuscript. This article is subject to HHMI’s Open Access to Publications policy. HHMI lab heads have previously granted a nonexclusive CC BY 4.0 license to the public and a sublicensable license to HHMI in their research articles. Pursuant to those licenses, the author-accepted manuscript of this article can be made freely available under a CC BY 4.0 license immediately upon publication.

## Authors’ contributions

Conceptualization, D.P., E.E.E.; Formal analysis, D.P.; Investigation, D.P., E.E.E.; Inversion genotyping, W.H., H.A., J.E., T.M.; Strand-seq data generation, A.D.S., P.H., J.O.K.; Computational support, W.T.H., A.N.R.; Assembly resources, B.P.; Writing, D.P., E.E.E., with input from all authors.

## Availability of data and materials

HPRC assemblies used in this study have been reported in (Liao et al. 2022). HGSVC assemblies used in this study have been reported in (Ebler et al. 2022). HPRC Strand-seq data: Data for eight samples (HG01123, HG01258, HG01358, HG01361, HG01891, HG02257, HG02486, HG02559) are available via ENA at accession number: PRJEB54100 (Porubsky, Vollger, et al. 2022). Additional Strand-seq data for samples HG01138 and HG01888, reported in this study, are available via ENA at accession numbers ERS13463611 and ERS13463612, respectively. Hi-C data for two samples (HG02011 and NA20847) presented in this study can be obtained at accession number: ERP123231 (Ebert et al. 2021).

## Declarations

**Competing interests**

E.E.E. is a scientific advisory board (SAB) member of Variant Bio, Inc. The following authors have previously disclosed a patent application (no. EP19169090) relevant to Strand-seq: J.O.K., T.M., and D.P. The other authors declare no competing interests.

## References

Aganezov, Sergey, Stephanie M. Yan, Daniela C. Soto, Melanie Kirsche, Samantha Zarate, Pavel Avdeyev, Dylan J. Taylor, et al. 2022. “A Complete Reference Genome Improves Analysis of Human Genetic Variation.” Science 376 (6588): eabl3533.

Altemose, Nicolas, Glennis A. Logsdon, Andrey V. Bzikadze, Pragya Sidhwani, Sasha A. Langley, Gina V. Caldas, Savannah J. Hoyt, et al. 2022. “Complete Genomic and Epigenetic Maps of Human Centromeres.” Science 376 (6588): eabl4178.

Antonacci, Francesca, Jeffrey M. Kidd, Tomas Marques-Bonet, Brian Teague, Mario Ventura, Santhosh Girirajan, Can Alkan, et al. 2010. “A Large and Complex Structural Polymorphism at 16p12.1 Underlies Microdeletion Disease Risk.” Nature Genetics 42 (9): 745–50.

Audano, Peter A., Arvis Sulovari, Tina A. Graves-Lindsay, Stuart Cantsilieris, Melanie Sorensen, Annemarie E. Welch, Max L. Dougherty, et al. 2019. “Characterizing the Major Structural Variant Alleles of the Human Genome.” Cell 176 (3): 663–75.e19.

Bragin, Eugene, Eleni A. Chatzimichali, Caroline F. Wright, Matthew E. Hurles, Helen V. Firth, A. Paul Bevan, and G. Jawahar Swaminathan. 2014. “DECIPHER: Database for the Interpretation of Phenotype-Linked Plausibly Pathogenic Sequence and Copy-Number Variation.” Nucleic Acids Research 42 (Database issue): D993–1000.

Cáceres, Mario, National Institutes of Health Intramural Sequencing Center Comparative Sequencing Program, Robert T. Sullivan, and James W. Thomas. 2007. “A Recurrent Inversion on the Eutherian X Chromosome.” Proceedings of the National Academy of Sciences of the United States of America 104 (47): 18571–76.

Chaisson, Mark J. P., Ashley D. Sanders, Xuefang Zhao, Ankit Malhotra, David Porubsky, Tobias Rausch, Eugene J. Gardner, et al. 2019. “Multi-Platform Discovery of Haplotype-Resolved Structural Variation in Human Genomes.” Nature Communications 10 (1): 1784.

Coe, Bradley P., Kali Witherspoon, Jill A. Rosenfeld, Bregje W. M. van Bon, Anneke T. Vulto-van Silfhout, Paolo Bosco, Kathryn L. Friend, et al. 2014. “Refining Analyses of Copy Number Variation Identifies Specific Genes Associated with Developmental Delay.” Nature Genetics 46 (10): 1063–71.

Cooper, Gregory M., Bradley P. Coe, Santhosh Girirajan, Jill A. Rosenfeld, Tiffany H. Vu, Carl Baker, Charles Williams, et al. 2011. “A Copy Number Variation Morbidity Map of Developmental Delay.” Nature Genetics 43 (9): 838–46.

Ebert, Peter, Peter A. Audano, Qihui Zhu, Bernardo Rodriguez-Martin, David Porubsky, Marc Jan Bonder, Arvis Sulovari, et al. 2021. “Haplotype-Resolved Diverse Human Genomes and Integrated Analysis of Structural Variation.” Science, February. https://doi.org/10.1126/science.abf7117.

Ebler, Jana, Peter Ebert, Wayne E. Clarke, Tobias Rausch, Peter A. Audano, Torsten Houwaart, Yafei Mao, et al. 2022. “Pangenome-Based Genome Inference Allows Efficient and Accurate Genotyping across a Wide Spectrum of Variant Classes.” Nature Genetics 54 (4): 518–25.

Eichler, E. E., H. A. Hammond, J. N. Macpherson, P. A. Ward, and D. L. Nelson. 1995. “Population Survey of the Human FMR1 CGG Repeat Substructure Suggests Biased Polarity for the Loss of AGG Interruptions.” Human Molecular Genetics 4 (12): 2199–2208.

Falconer, Ester, Mark Hills, Ulrike Naumann, Steven S. S. Poon, Elizabeth A. Chavez, Ashley D. Sanders, Yongjun Zhao, Martin Hirst, and Peter M. Lansdorp. 2012. “DNA Template Strand Sequencing of Single-Cells Maps Genomic Rearrangements at High Resolution.” Nature Methods 9 (11): 1107–12.

Gebhardt, Gabriel Stefan, Koenraad Devriendt, Reinhilde Thoelen, Ann Swillen, Elly Pijkels, Jean-Pierre Fryns, Joris R. Vermeesch, and Marc Gewillig. 2003. “No Evidence for a Parental Inversion Polymorphism Predisposing to Rearrangements at 22q11.2 in the DiGeorge/Velocardiofacial Syndrome.” European Journal of Human Genetics: EJHG 11 (2): 109–11.

Gel, Bernat, Anna Díez-Villanueva, Eduard Serra, Marcus Buschbeck, Miguel A. Peinado, and Roberto Malinverni. 2016. “regioneR: An R/Bioconductor Package for the Association Analysis of Genomic Regions Based on Permutation Tests.” Bioinformatics 32 (2): 289–91.

Hanlon, Vincent C. T., Peter M. Lansdorp, and Victor Guryev. 2022. “A Survey of Current Methods to Detect and Genotype Inversions.” Human Mutation, September. https://doi.org/10.1002/humu.24458.

Itsara, Andy, Gregory M. Cooper, Carl Baker, Santhosh Girirajan, Jun Li, Devin Absher, Ronald M. Krauss, et al. 2009. “Population Analysis of Large Copy Number Variants and Hotspots of Human Genetic Disease.” American Journal of Human Genetics 84 (2): 148–61.

Jarvis, Erich D., Giulio Formenti, Arang Rhie, Andrea Guarracino, Chentao Yang, Jonathan Wood, Alan Tracey, et al. 2022. “Automated Assembly of High-Quality Diploid Human Reference Genomes.” bioRxiv. https://doi.org/10.1101/2022.03.06.483034.

Kidd, Jeffrey M., Gregory M. Cooper, William F. Donahue, Hillary S. Hayden, Nick Sampas, Tina Graves, Nancy Hansen, et al. 2008. “Mapping and Sequencing of Structural Variation from Eight Human Genomes.” Nature 453 (7191): 56–64.

Kidd, Jeffrey M., Tina Graves, Tera L. Newman, Robert Fulton, Hillary S. Hayden, Maika Malig, Joelle Kallicki, Rajinder Kaul, Richard K. Wilson, and Evan E. Eichler. 2010. “A Human Genome Structural Variation Sequencing Resource Reveals Insights into Mutational Mechanisms.” Cell 143 (5): 837–47.

Kidd, Jeffrey M., Nick Sampas, Francesca Antonacci, Tina Graves, Robert Fulton, Hillary S. Hayden, Can Alkan, et al. 2010. “Characterization of Missing Human Genome Sequences and Copy-Number Polymorphic Insertions.” Nature Methods 7 (5): 365–71.

Kong, Augustine, Gudmar Thorleifsson, Daniel F. Gudbjartsson, Gisli Masson, Asgeir Sigurdsson, Aslaug Jonasdottir, G. Bragi Walters, et al. 2010. “Fine-Scale Recombination Rate Differences between Sexes, Populations and Individuals.” Nature 467 (7319): 1099–1103.

Koolen, David A., Lisenka E. L. Vissers, Rolph Pfundt, Nicole de Leeuw, Samantha J. L. Knight, Regina Regan, R. Frank Kooy, et al. 2006. “A New Chromosome 17q21.31 Microdeletion Syndrome Associated with a Common Inversion Polymorphism.” Nature Genetics. https://doi.org/10.1038/ng1853.

Liao, Wen-Wei, Mobin Asri, Jana Ebler, Daniel Doerr, Marina Haukness, Glenn Hickey, Shuangjia Lu, et al. 2022. “A Draft Human Pangenome Reference.” bioRxiv. https://doi.org/10.1101/2022.07.09.499321.

Li, Heng, and Richard Durbin. 2010. “Fast and Accurate Long-Read Alignment with Burrows–Wheeler Transform.” Bioinformatics 26 (5): 589–95.

Lun, Aaron T. L., and Gordon K. Smyth. 2015. “diffHic: A Bioconductor Package to Detect Differential Genomic Interactions in Hi-C Data.” BMC Bioinformatics 16 (August): 258.

Maggiolini, Flavia A. M., Stuart Cantsilieris, Pietro D’Addabbo, Michele Manganelli, Bradley P. Coe, Beth L. Dumont, Ashley D. Sanders, et al. 2019. “Genomic Inversions and GOLGA Core Duplicons Underlie Disease Instability at the 15q25 Locus.” PLoS Genetics 15 (3): e1008075.

Mohajeri, Kiana, Stuart Cantsilieris, John Huddleston, Bradley J. Nelson, Bradley P. Coe, Catarina D. Campbell, Carl Baker, et al. 2016. “Interchromosomal Core Duplicons Drive Both Evolutionary Instability and Disease Susceptibility of the Chromosome 8p23.1 Region.” Genome Research 26 (11): 1453–67.

Nurk, Sergey, Sergey Koren, Arang Rhie, Mikko Rautiainen, Andrey V. Bzikadze, Alla Mikheenko, Mitchell R. Vollger, et al. 2021. “The Complete Sequence of a Human Genome.” bioRxiv. https://doi.org/10.1101/2021.05.26.445798.

Osborne, L. R., M. Li, B. Pober, D. Chitayat, J. Bodurtha, A. Mandel, T. Costa, et al. 2001. “A 1.5 Million-Base Pair Inversion Polymorphism in Families with Williams-Beuren Syndrome.” Nature Genetics 29 (3): 321–25.

Osman, I., H. Scher, G. Dalbagni, V. Reuter, Z. F. Zhang, and C. Cordon-Cardo. 1997. “Chromosome 16 in Primary Prostate Cancer: A Microsatellite Analysis.” International Journal of Cancer. Journal International Du Cancer 71 (4): 580–84.

Porubsky, David, Peter Ebert, Peter A. Audano, Mitchell R. Vollger, William T. Harvey, Pierre Marijon, Jana Ebler, et al. 2021. “Fully Phased Human Genome Assembly without Parental Data Using Single-Cell Strand Sequencing and Long Reads.” Nature Biotechnology 39 (3): 302–8.

Porubsky, David, Wolfram Höps, Hufsah Ashraf, Pinghsun Hsieh, Bernardo Rodriguez-Martin, Feyza Yilmaz, Jana Ebler, et al. 2022. “Recurrent Inversion Polymorphisms in Humans Associate with Genetic Instability and Genomic Disorders.” Cell, May. https://doi.org/10.1016/j.cell.2022.04.017.

Porubsky, David, Ashley D. Sanders, Wolfram Höps, Pinghsun Hsieh, Arvis Sulovari, Ruiyang Li, Ludovica Mercuri, et al. 2020. “Recurrent Inversion Toggling and Great Ape Genome Evolution.” Nature Genetics, June. https://doi.org/10.1038/s41588-020-0646-x.

Porubsky, David, Mitchell R. Vollger, William T. Harvey, Allison N. Rozanski, Peter Ebert, Glenn Hickey, Patrick Hasenfeld, et al. 2022. “Gaps and Complex Structurally Variant Loci in Phased Genome Assemblies.” bioRxiv. https://doi.org/10.1101/2022.07.06.498874.

Puig, Marta, Jon Lerga-Jaso, Carla Giner-Delgado, Sarai Pacheco, David Izquierdo, Alejandra Delprat, Magdalena Gayà-Vidal, Jack F. Regan, George Karlin-Neumann, and Mario Cáceres. 2020. “Determining the Impact of Uncharacterized Inversions in the Human Genome by Droplet Digital PCR.” Genome Research 30 (5): 724–35.

Rautiainen, Mikko, Sergey Nurk, Brian P. Walenz, Glennis A. Logsdon, David Porubsky, Arang Rhie, Evan E. Eichler, Adam M. Phillippy, and Sergey Koren. 2022. “Verkko: Telomere-to-Telomere Assembly of Diploid Chromosomes.” bioRxiv. https://doi.org/10.1101/2022.06.24.497523.

Sanders, Ashley D., Ester Falconer, Mark Hills, Diana C. J. Spierings, and Peter M. Lansdorp. 2017. “Single-Cell Template Strand Sequencing by Strand-Seq Enables the Characterization of Individual Homologs.” Nature Protocols 12 (6): 1151–76.

Sanders, Ashley D., Mark Hills, David Porubský, Victor Guryev, Ester Falconer, and Peter M. Lansdorp. 2016. “Characterizing Polymorphic Inversions in Human Genomes by Single-Cell Sequencing.” Genome Research 26 (11): 1575–87.

Sanders, Ashley D., Sascha Meiers, Maryam Ghareghani, David Porubsky, Hyobin Jeong, M. Alexandra C. C. van Vliet, Tobias Rausch, et al. 2020. “Single-Cell Analysis of Structural Variations and Complex Rearrangements with Tri-Channel Processing.” Nature Biotechnology 38 (3): 343–54.

She, Xinwei, Zhaoshi Jiang, Royden A. Clark, Ge Liu, Ze Cheng, Eray Tuzun, Deanna M. Church, Granger Sutton, Aaron L. Halpern, and Evan E. Eichler. 2004. “Shotgun Sequence Assembly and Recent Segmental Duplications within the Human Genome.” Nature 431 (7011): 927–30.

Steinberg, Karyn Meltz, Francesca Antonacci, Peter H. Sudmant, Jeffrey M. Kidd, Catarina D. Campbell, Laura Vives, Maika Malig, et al. 2012. “Structural Diversity and African Origin of the 17q21.31 Inversion Polymorphism.” Nature Genetics. https://doi.org/10.1038/ng.2335.

Sturtevant, A. H. 1917. “Genetic Factors Affecting the Strength of Linkage in Drosophila.” Proceedings of the National Academy of Sciences of the United States of America 3 (9): 555–58.

Tarasov, Artem, Albert J. Vilella, Edwin Cuppen, Isaac J. Nijman, and Pjotr Prins. 2015. “Sambamba: Fast Processing of NGS Alignment Formats.” Bioinformatics 31 (12): 2032–34.

Vergés, Laia, Francesca Vidal, Esther Geán, Alexandra Alemany-Schmidt, Maria Oliver-Bonet, and Joan Blanco. 2017. “An Exploratory Study of Predisposing Genetic Factors for DiGeorge/velocardiofacial Syndrome.” Scientific Reports 7 (January): 40031.

Vollger, Mitchell R., Xavi Guitart, Philip C. Dishuck, Ludovica Mercuri, William T. Harvey, Ariel Gershman, Mark Diekhans, et al. 2022. “Segmental Duplications and Their Variation in a Complete Human Genome.” Science 376 (6588): eabj6965.

Wang, Ting, Lucinda Antonacci-Fulton, Kerstin Howe, Heather A. Lawson, Julian K. Lucas, Adam M. Phillippy, Alice B. Popejoy, et al. 2022. “The Human Pangenome Project: A Global Resource to Map Genomic Diversity.” Nature 604 (7906): 437–46.

Zody, Michael C., Zhaoshi Jiang, Hon-Chung Fung, Francesca Antonacci, Ladeana W. Hillier, Maria Francesca Cardone, Tina A. Graves, et al. 2008. “Evolutionary Toggling of the MAPT 17q21.31 Inversion Region.” Nature Genetics 40 (9): 1076–83.

