## Supplemental Material for "Inversion polymorphism in a complete human genome assembly"

#### SUPPLEMENTAL FIGURES (S1-S18):

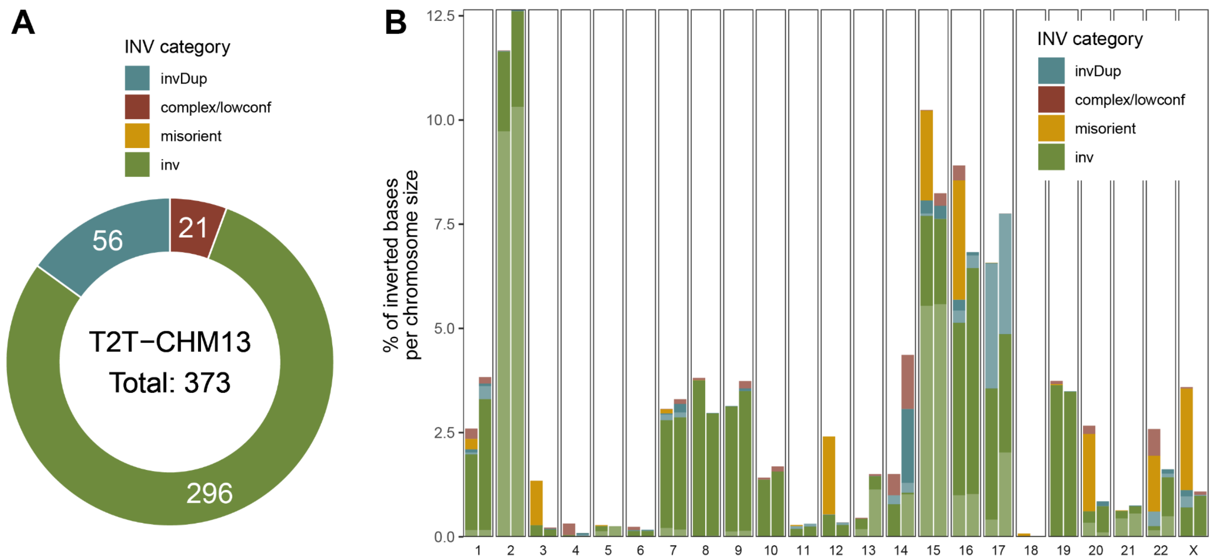


**Figure S1: T2T-CHM13 inversion callset summary and comparison to GRCh38 (n=373).**
**A**) A donut plot showing the total counts of inversion classes defined based on Arbigent and PAV genotypes. **B**) A barplot showing the percentage of inverted bases per inversion category (misorient - misorientation, inv - balanced inversion, invDup - inverted duplication and complex/lowconf - structurally complex region or low-confidence call) given the chromosome size. For each chromosome, the left- and right-side bars represent the fraction of inverted bases in GRCh38 and T2T-CHM13, respectively. Lighter color highlights bases being inverted only in a single sample (see light green color for chromosome 2 contributed by a single pericentromeric inversion ~23 Mbp in size in sample NA19650).

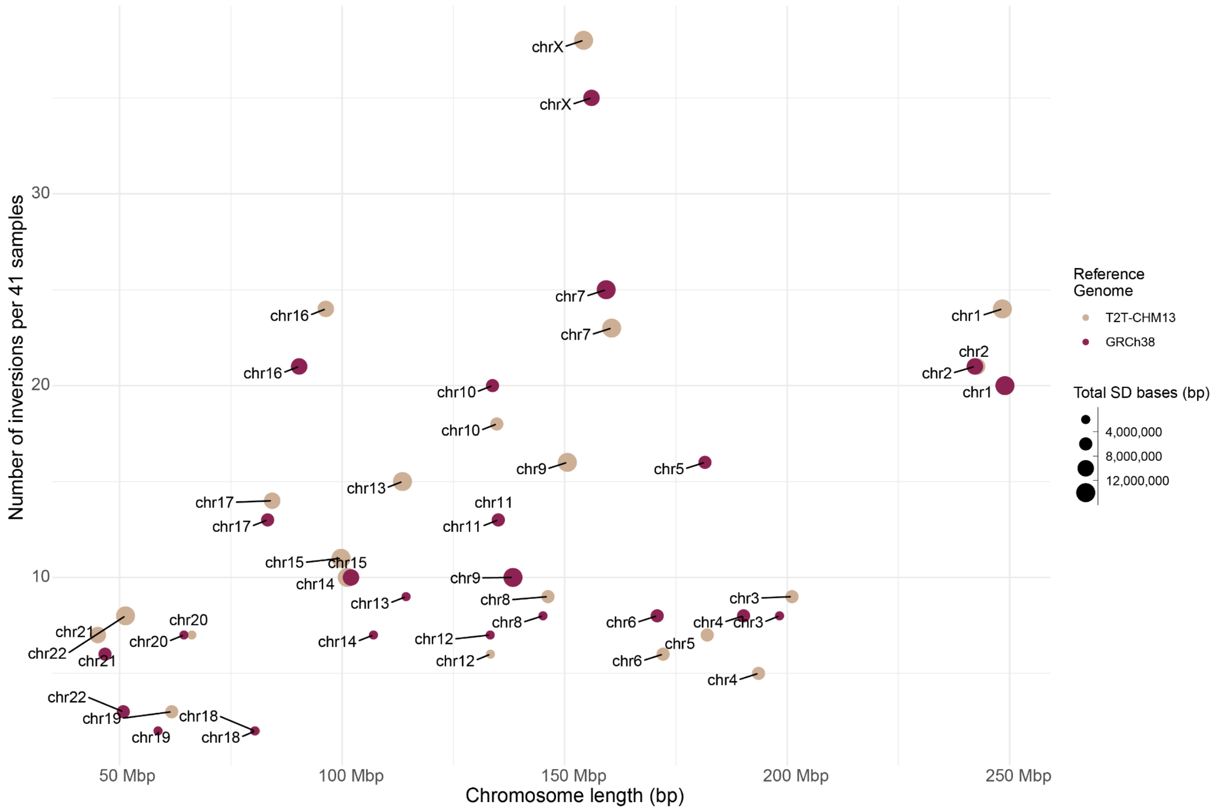


**Figure S2: Differences between GRCh38 and T2T-CHM13 callsets. (join dots only one chr label)**

A scatterplot shows the total number of balanced inversions detected per chromosome (y-axis) given the chromosome length (x-axis) separately per GRCh38 (beige) and T2T-CHM13 (purple) inversion callset. Size of each dot represents the total number of SD bases reported for a given chromosome and given reference.

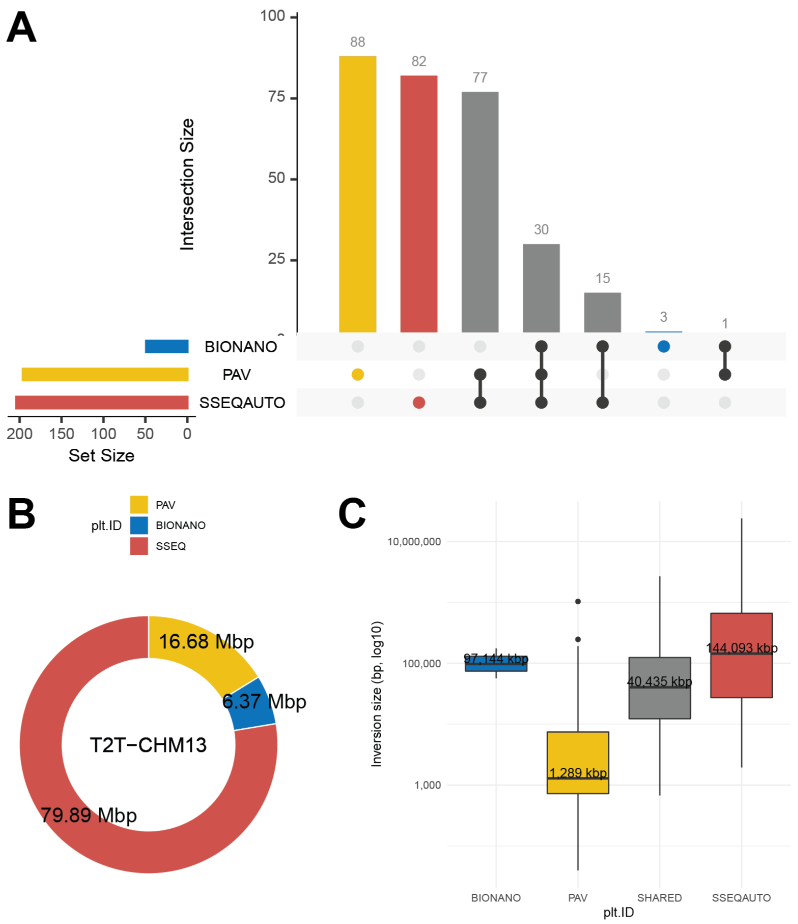


**Figure S3: Inversion callset summary with respect to CHM13 reference.**
**A**) An upset plot showing the total number of inversions uniquely detected by each technology and those detected by two and more. **B**) A donut plot showing the number of inverted kilobases contributed separately by each technology to the final callset. **C**) A boxplot showing the size distribution of inversions uniquely detected by each technology and those detected by two and more (SHARED).

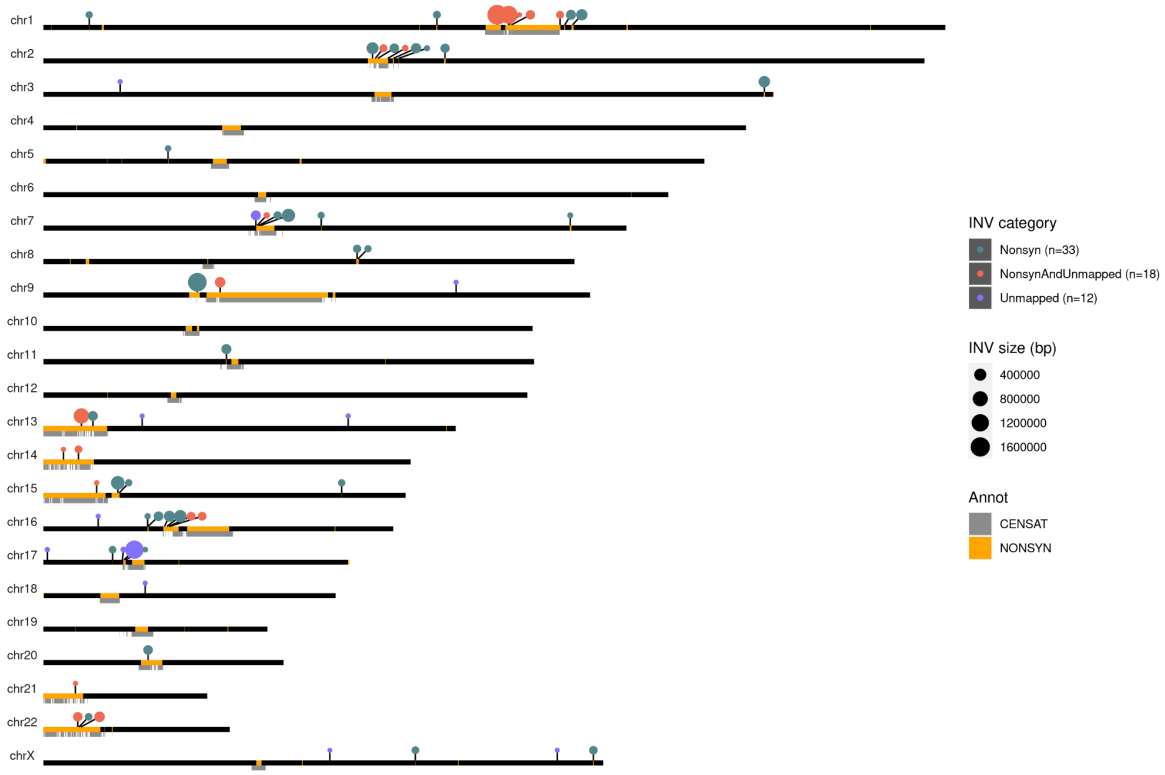


**Figure S4: Non-syntenic and likely novel sites in T2T-CHM13 inversion calls.**
An ideogram showing the position and size (dot size) of all balanced inversions (n=63) that either fall within (>=90% reciprocal overlap) non-syntenic regions between GRCh38 and T2T-CHM13 ('Nonsyn') or failed to map to GRCh38 reference ('Unmapped') (**Methods, Table S2**). Red dots point to regions whose sequence failed both to map to GRCh38 reference and fall within nonsyntenic regions (n=18).

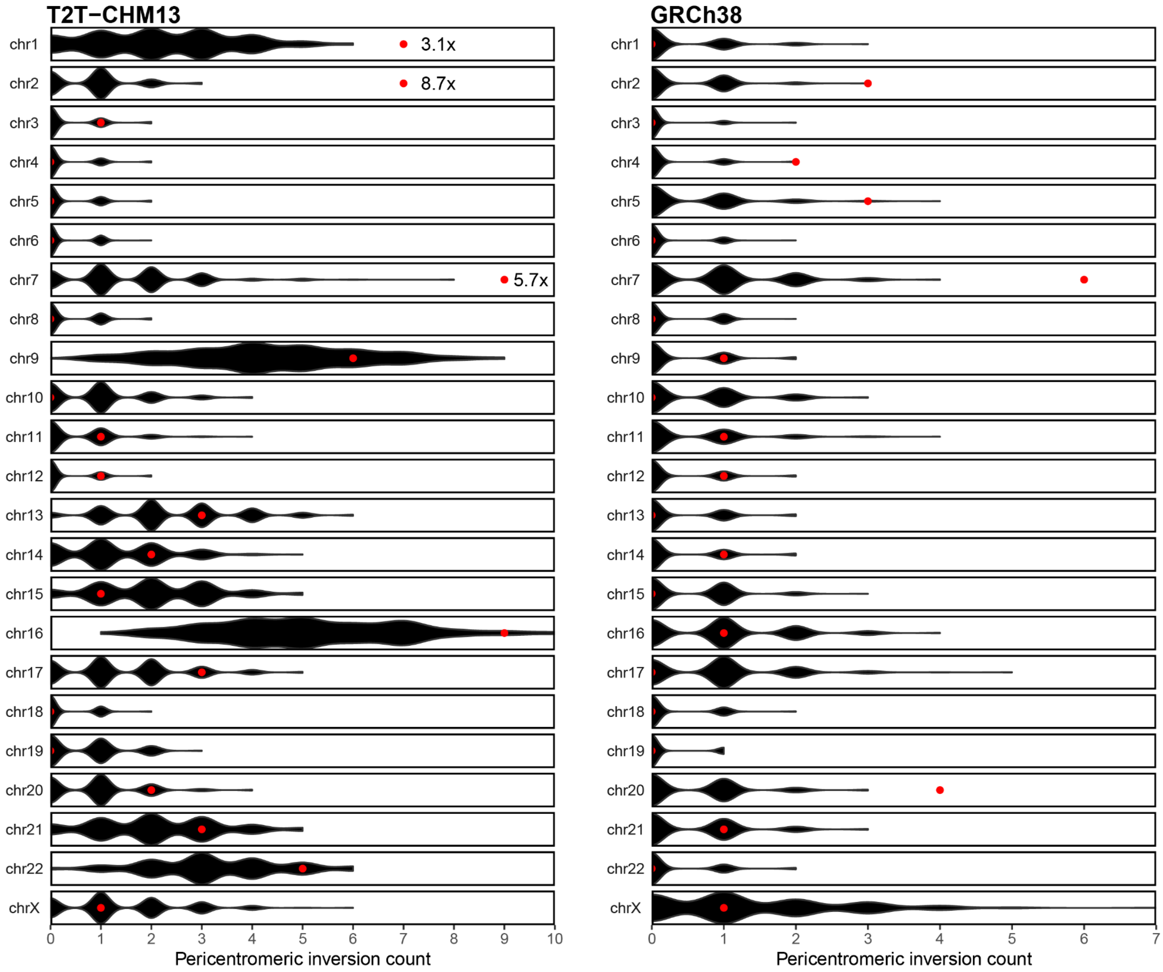


**Figure S5: Enrichment of inversion in pericentromeric regions.**
Permutation analysis of inversion falling within the pericentromeric region of each chromosome. Permuted counts of pericentromeric inversions are shown as black violin plots. Observed values are shown as red dots. Enrichment analysis is reported separately with respect to T2T-CHM13 (left) and GRCh38 (right) (**Methods**). For T2T-CHM13 reference we highlight fold-enrichment values for chromosomes that reached significance after Bonferroni correction (**Table S3**).

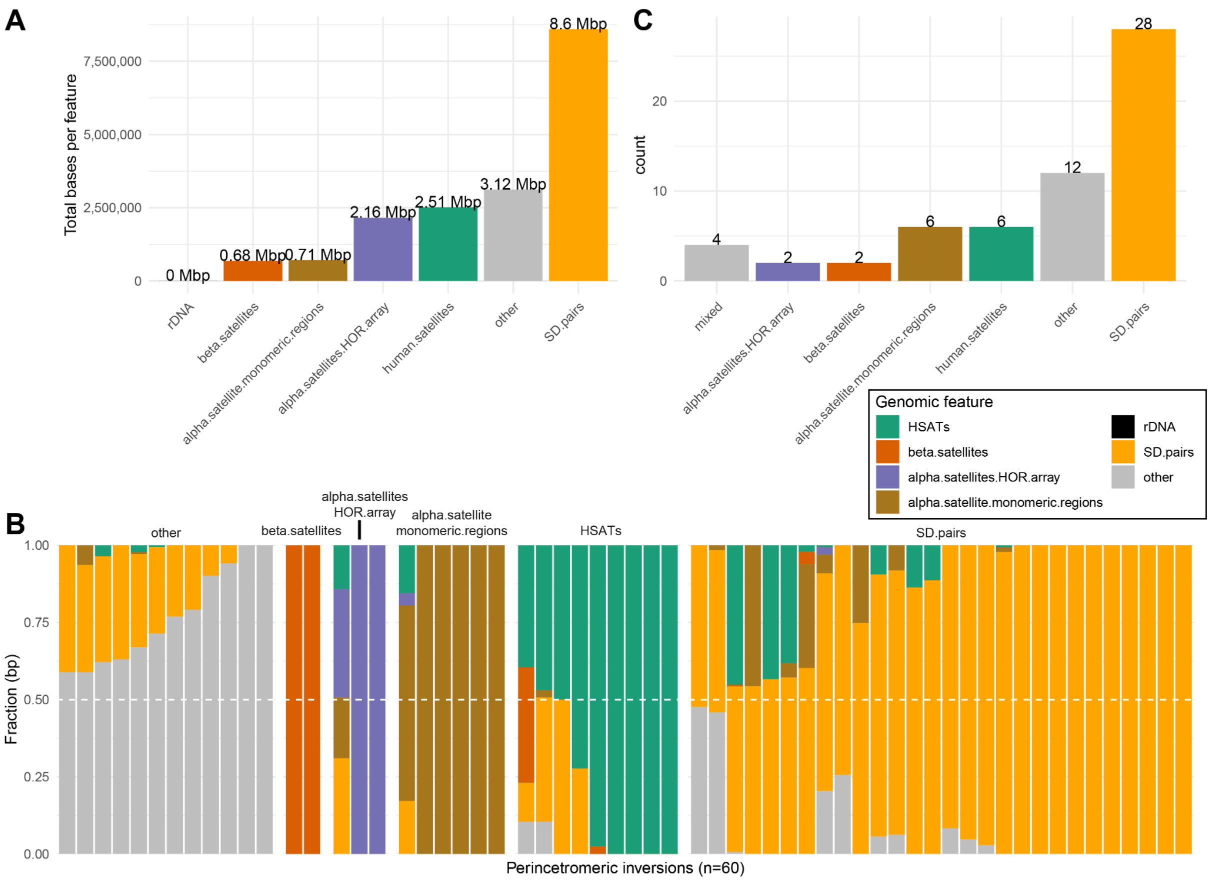


#### Figure S6: Sequence composition of inversions from pericentromeric regions.

**A**) The total number of base pairs of various genomic features (such as various classes of human satellites, 'SD.pairs' - intrachromosomal pairs of SDs no further than 5 Mbp apart and ‘other’ - none of these features) overlapping pericentromeric inversions (n=60). **B**) Proportion of genomic features assigned to each brnn region based on the number of ‘burned’ haplotypes within each brnn region. **C**) An assignment of each pericentromeric inversion to a single feature based on the majority overlap (proportion of the given feature >0.5) or are labeled as ‘mixed’ if no feature is >0.5.

**NOTE**: In this analysis we excluded the large pericentromeric inversion on chromosome 2 that is ~23 Mbp in size to prevent our results being skewed by including such a large genomic region.

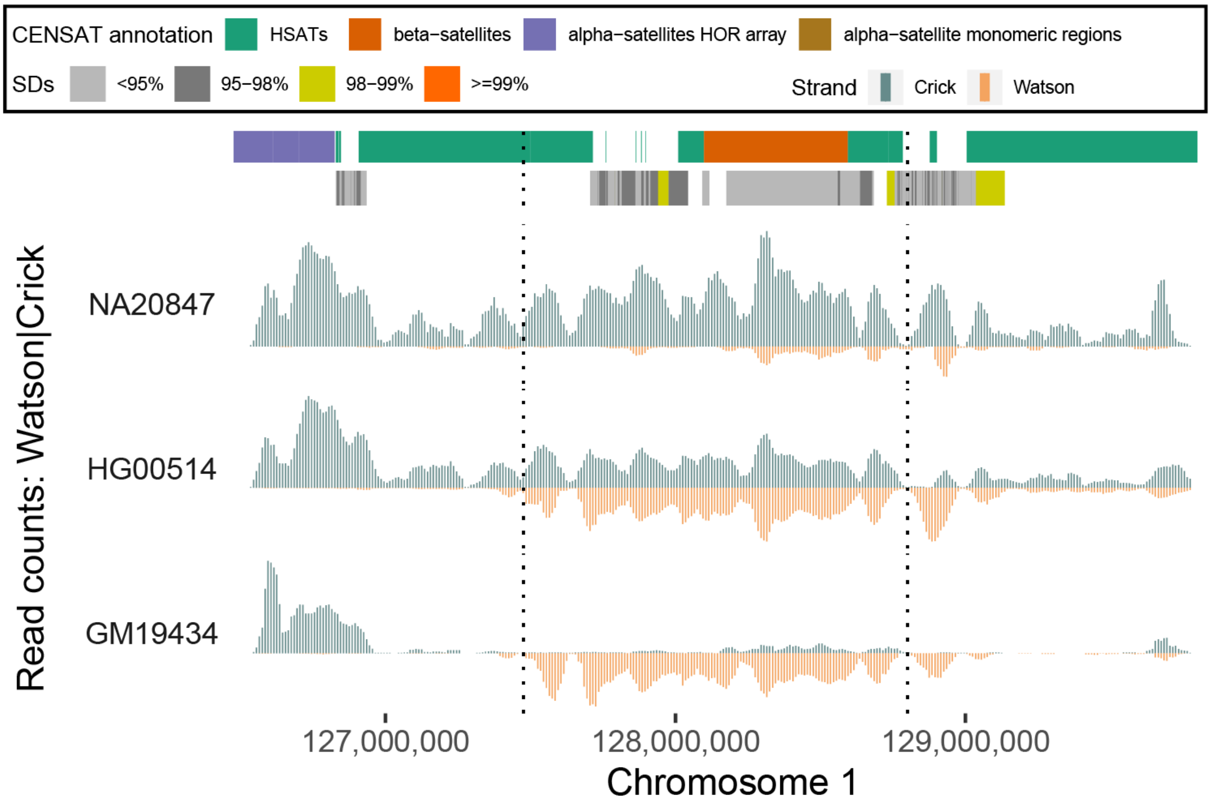


**Figure S7: Novel pericentromeric inversion on chromosome 1.**
A zoomed-in plot on novel pericentromeric inversion on chromosome 1 (highlighted by dotted lines) presented in Fig. 1C. The read-coverage profiles of Strand-seq data over a chromosome 1 centromeric region summarized as binned (bin size: 50 kbp step size: 10 kbp) read counts represented as bars above (teal; Crick read counts) and below (orange; Watson read counts) the midline. Dotted lines highlight the novel centromeric inversion detected on chromosome 1 only with respect to T2T-CHM13. In this region equal coverage of Watson and Crick count represents a heterozygous inversion as only one homologue is inverted with respect to the reference while reads aligned only in Watson orientation represents a homozygous inversion. Above is a centromere and SD annotation.

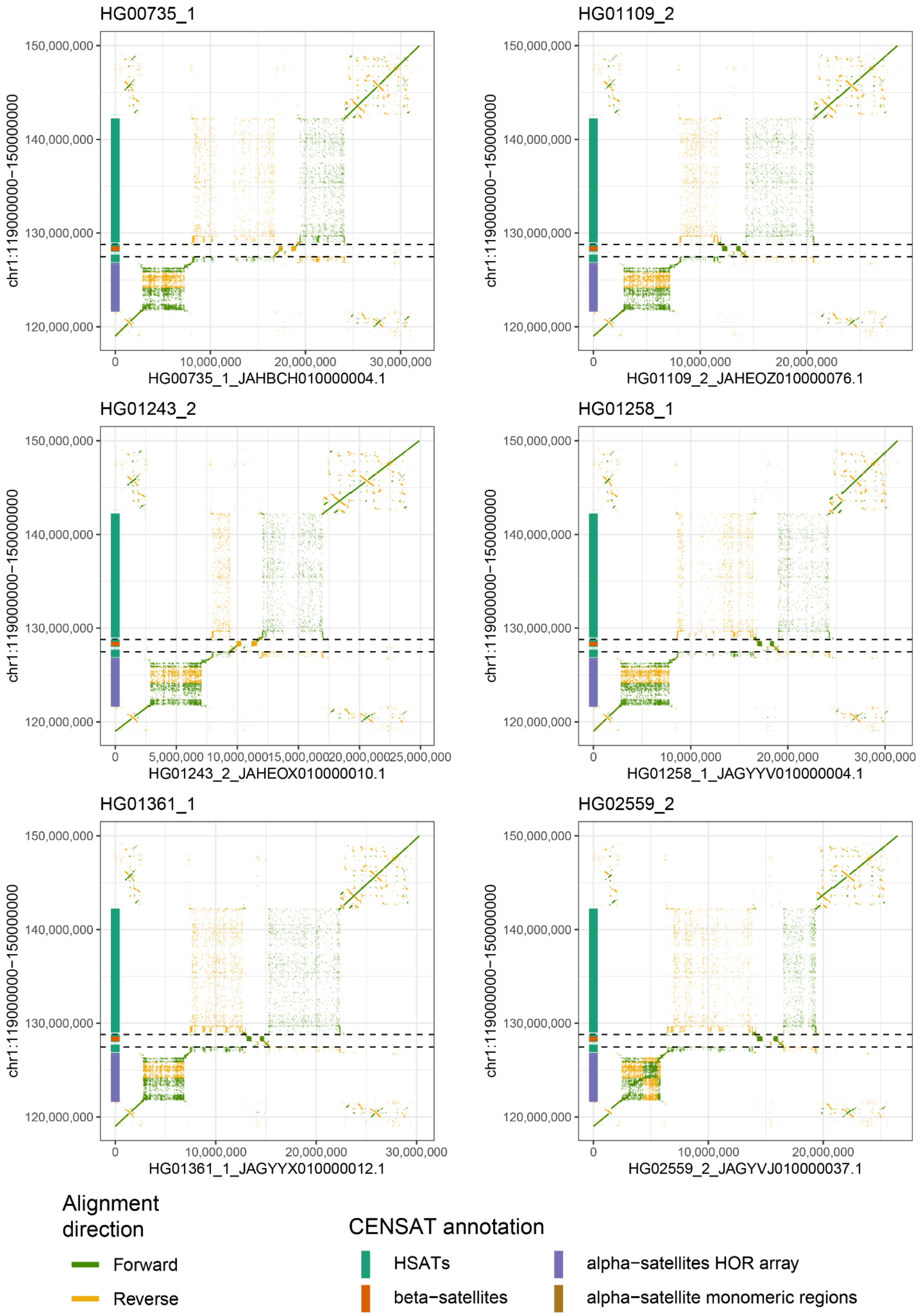


**Figure S8: Complete assemblies of chromosome 1 centromeric region.**
Dotplots showing the alignment directionality (yellow - reverse, green - forward) between complete assemblies of chromosome 1 centromere (x-axis) against T2T-CHM13 reference (y-axis). T2T-CHM13 centromere annotation is shown as colored boxes on the y-axis.

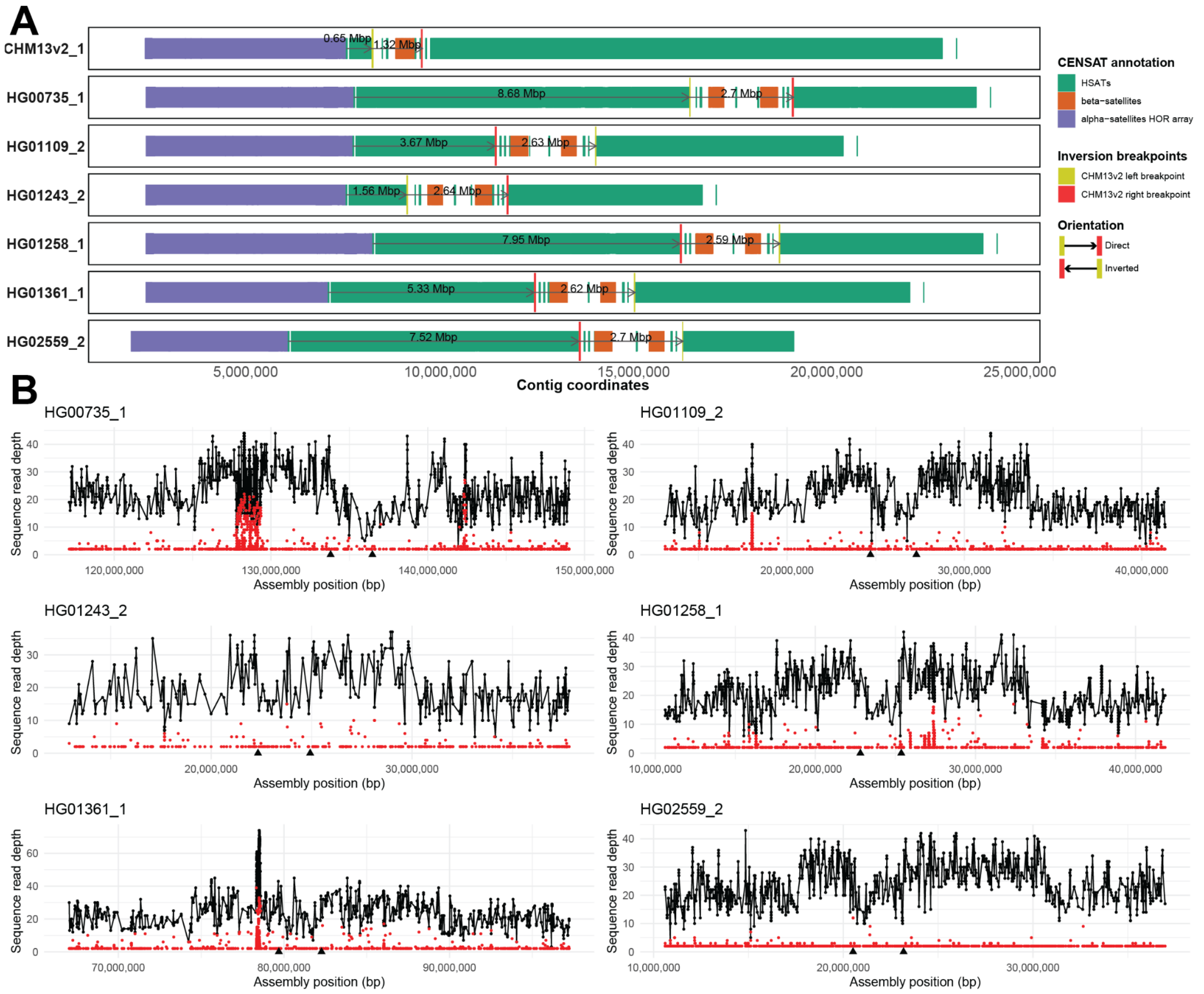


**Figure S9: Relative position of alpha satellite array and novel pericentromeric inversion on chromosome 1.**

**A)** RepeatMasker annotation of centromeric repeats for six complete assemblies of chromosome 1 centromere along with T2T-CHM13 reference (CHM13v2_1). We highlight alpha satellites (purple), beta satellites (orange) and human satellites (green). Mapped left (yellow) and right (red) inversion breakpoints are highlighted as vertical bars (**Methods**). Distances between boundaries of alpha satellite repeats and left-most inversion breakpoints are shown as an arrow with a distance in Mbp. Similarly, we show the distance between left and right inversion breakpoints. **B**) NucFreq validation of assembled centromeric regions presented in A). Black dots show read depth for the most common base at a given position while red dots show the second most common base. Regions where we observe high depth of the second most abundant base (red) are likely assembly collapses. Predicted inversion breakpoints are marked as black arrowheads at the bottom of each plot.

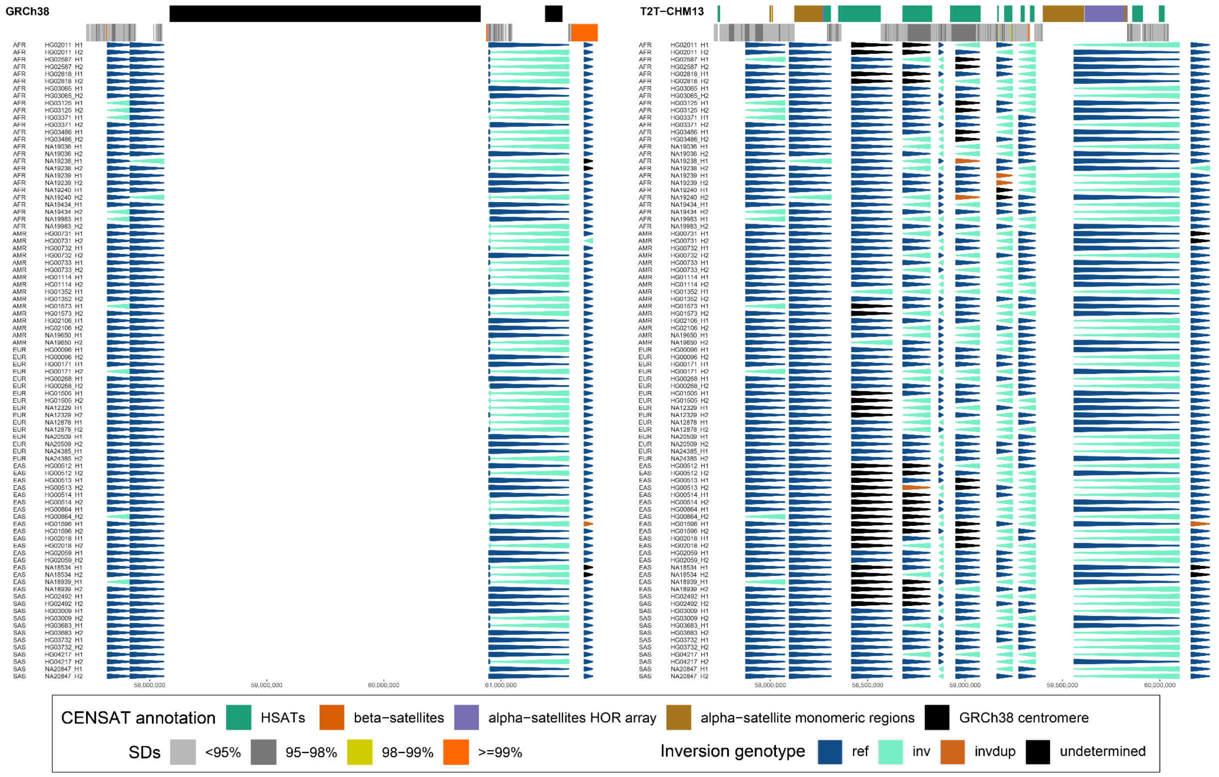


**Figure S10: Inversion phasing at pericentromeric region of chromosome 7.**
An arrowhead plot showing the inverted status of each defined region reported as colored arrowheads (dark blue - direct, bright blue - inverted, see the legend) for corresponding regions with respect to GRCh38 (left) and T2T-CHM13 (right). Gray rectangle in the middle highlights the positions of chromosome 7 centromere in GRCh38.

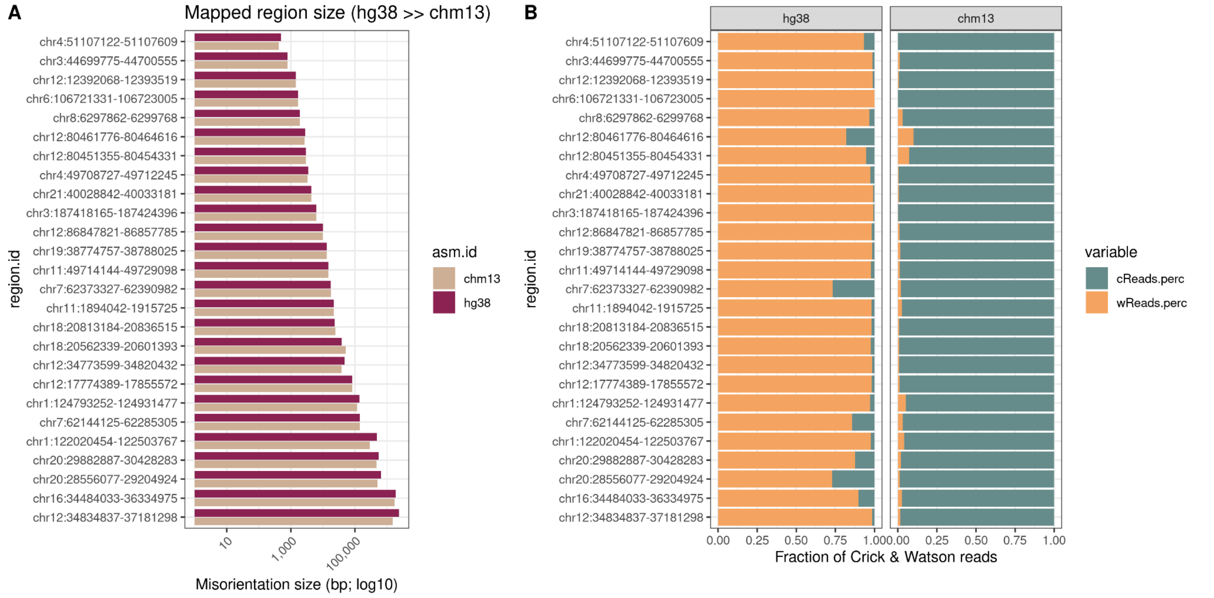


**Figure S11: Evaluation of putative misorients in GRCh38 with respect to T2T-CHM13.**
**A**) Distribution of region sizes of 28 putative misorients in GRCh38 (green) and their respective sizes after mapping onto the T2T-CHM13 reference genome (**Methods**). **B**) Shows fraction of Watson (minus; orange; wReads) and Crick (plus; teal; cReads) reads mapped to each region separately for reads mapped to GRCh38 and T2T-CHM13 reference genome. Read counts are concatenated across all unrelated individuals (n=41) reported in this study.

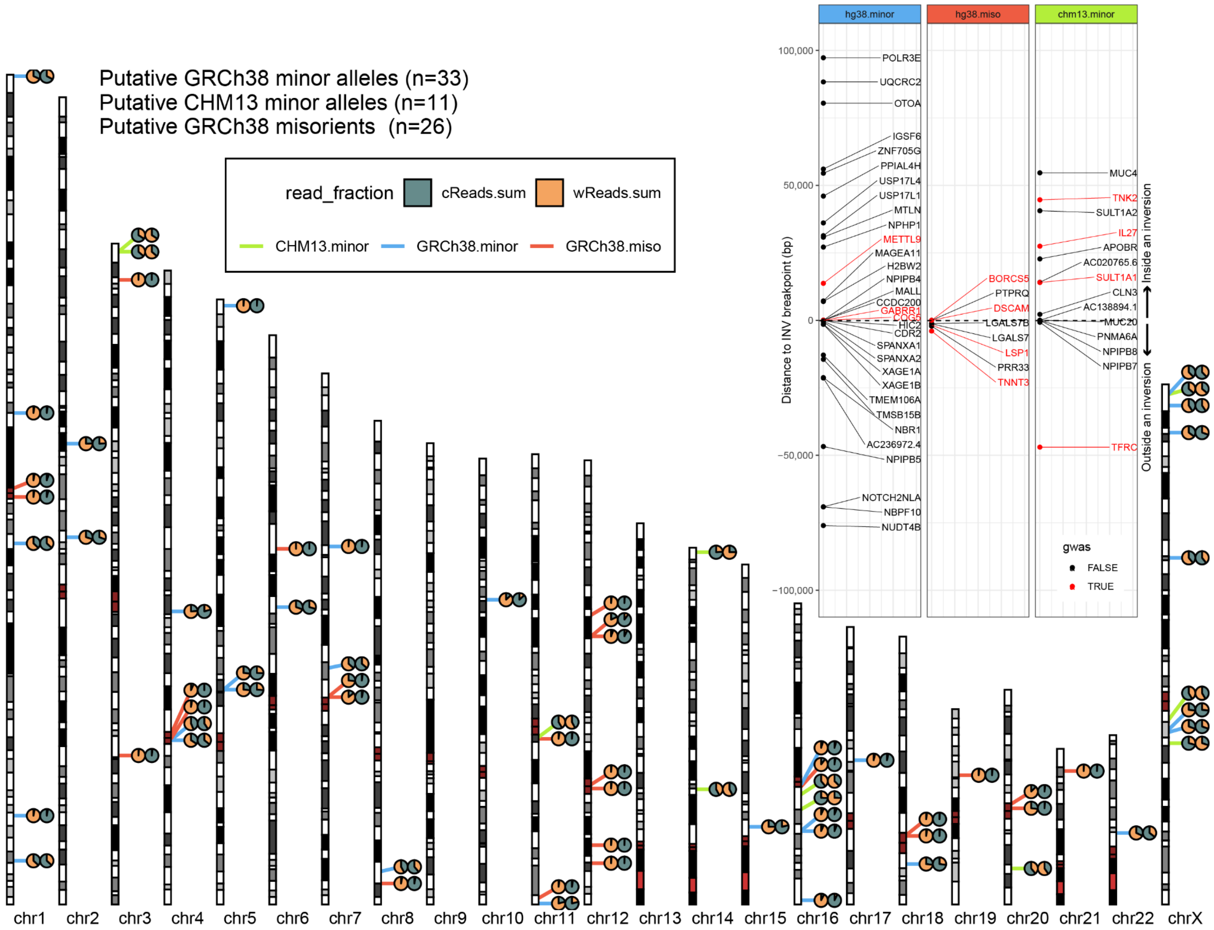


**Figure S12: Evaluation of inversion differences between GRCh38 and T2T-CHM13 reference.** A GRCh38 ideogram showing the fraction of Watson (orange; minus) and Crick (teal; plus) reads aligned to both GRCh38 (left side pie) and T2T-CHM13 (right side pie) reference for a selected number of regions. Strand-seq read counts are summarized across all unrelated individuals (n=41) from this study. Position of putative minor alleles in GRCh38 (n=33, blue lines) reference with respect to T2T-CHM13. Putative misorientations in GRCh38 (n=26) evaluated with respect to T2T-CHM13 are highlighted by red lines. Putative minor alleles in T2T-CHM13 (n=11) predicted with respect to GRCh38 are highlighted by green lines. **Inset:** Shows positions of protein-coding genes that reside within 100 kbp distance from GRCh38 misorientation (n=8), GRCh38 minor alleles (n=37) or T2T-CHM13 minor alleles (n=14). Gene names colored in red have been previously reported as part of genome-wide association studies (GWAS).

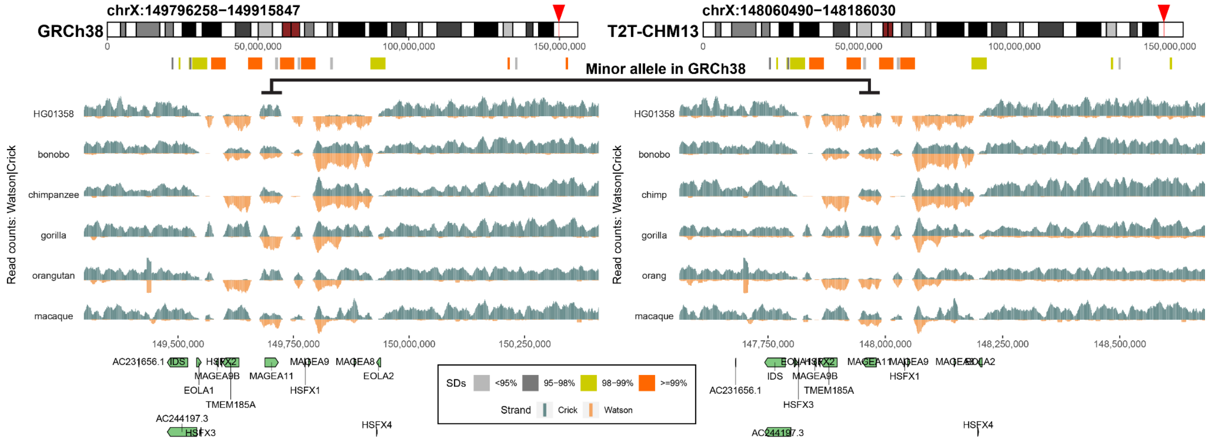


### Figure S13: Structural differences at Xq28 between GRCh38 and T2T-CHM13.

Strand-seq read-coverage profiles over a Xq28 region summarized as binned (bin size: 10 kbp step size: 1 kbp) read counts represented as bars above (teal; Crick read counts) and below (orange; Watson read counts) the midline. An equal coverage of Watson and Crick count represents a heterozygous inversion as only one homologue is inverted with respect to the reference while reads aligned only in Watson orientation represents a homozygous inversion. There is a novel inversion in sample HG01358 with respect to T2T-CHM13. A horizontal line shows a region where there is a minor allele in GRCh38.

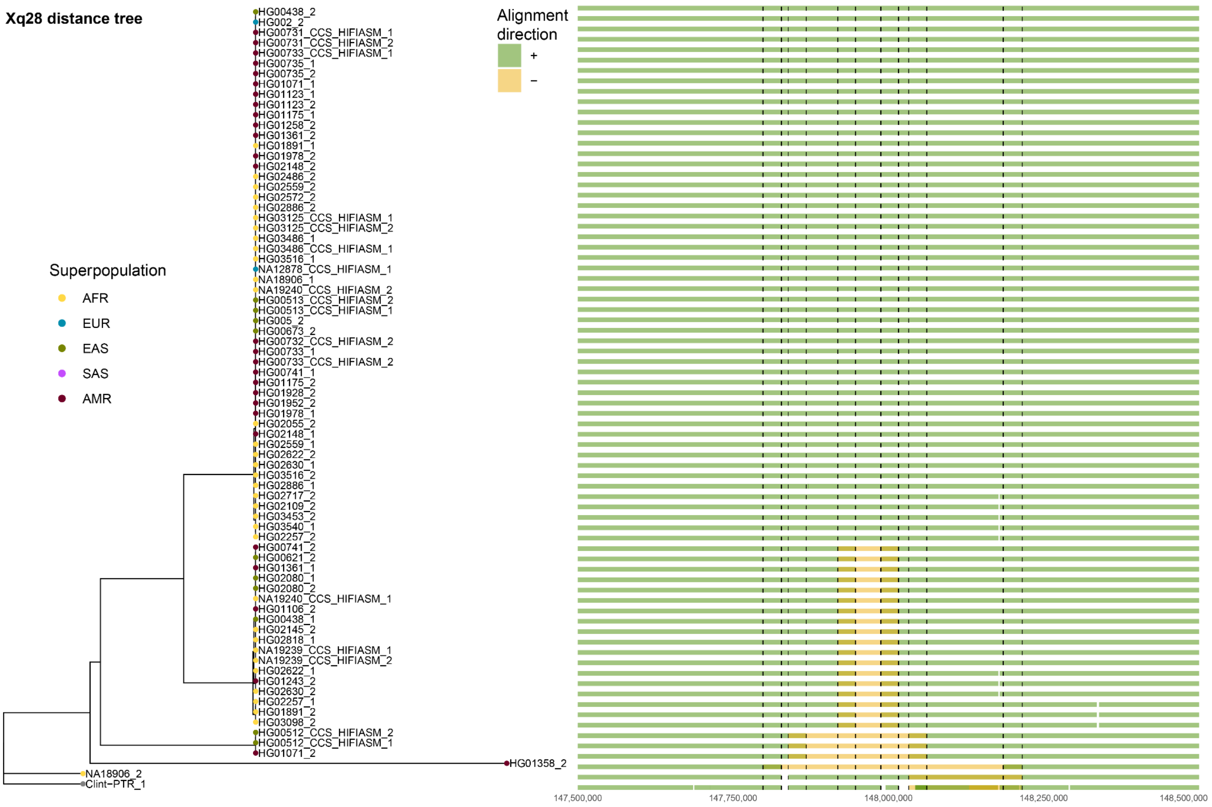


### Figure S14: Diverse structural haplotypes at Xq28 region.

Left: An UPGMA tree grouping complete assemblies (n=76) of Xq28 region into structurally similar groups based on their alignment to T2T-CHM13 (**Methods**). Superpopulation of origin for each sample is marked by colored dots. Right: Visualization of alignment directionality (plus - green, minus - orange) of each assembly with respect to T2T-CHM13. Positions of SD blocks in the Xq28 region are highlighted by vertical dotted lines. Each alignment is plotted with 0.5 level of transparency such that overlapping alignments are visible as boxes with a darker color or a mixed green and orange color.

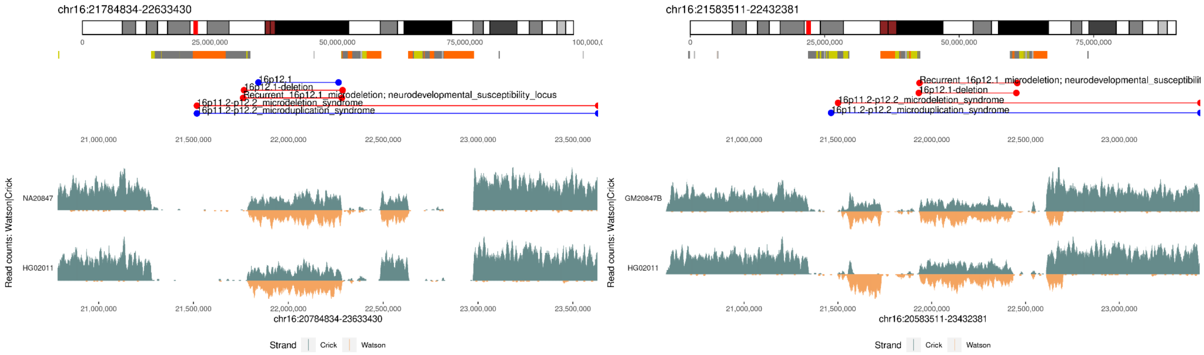


**Figure S15: Structural differences at 16p12.2 between GRCh38 and T2T-CHM13.**
Read-coverage profiles of Strand-seq data for a selected region on chromosome 16 with Strand-seq reads mapped separately to GRCh38 (left plot) and T2T-CHM13 (right plot) reference. Strand-seq reads are summarized as binned (bin size: 10 kbp, step size: 1 kbp) read counts represented as bars above (teal; Crick read counts) and below (orange; Watson read counts) midline. Region with roughly equal coverage of Watson and Crick count represents a heterozygous inversion as only one homologue is inverted with respect to the reference while region with reads aligned only in Watson orientation represents a homozygous inversion. Each inverted region is highlighted on chromosome-specific ideogram by a red rectangle.

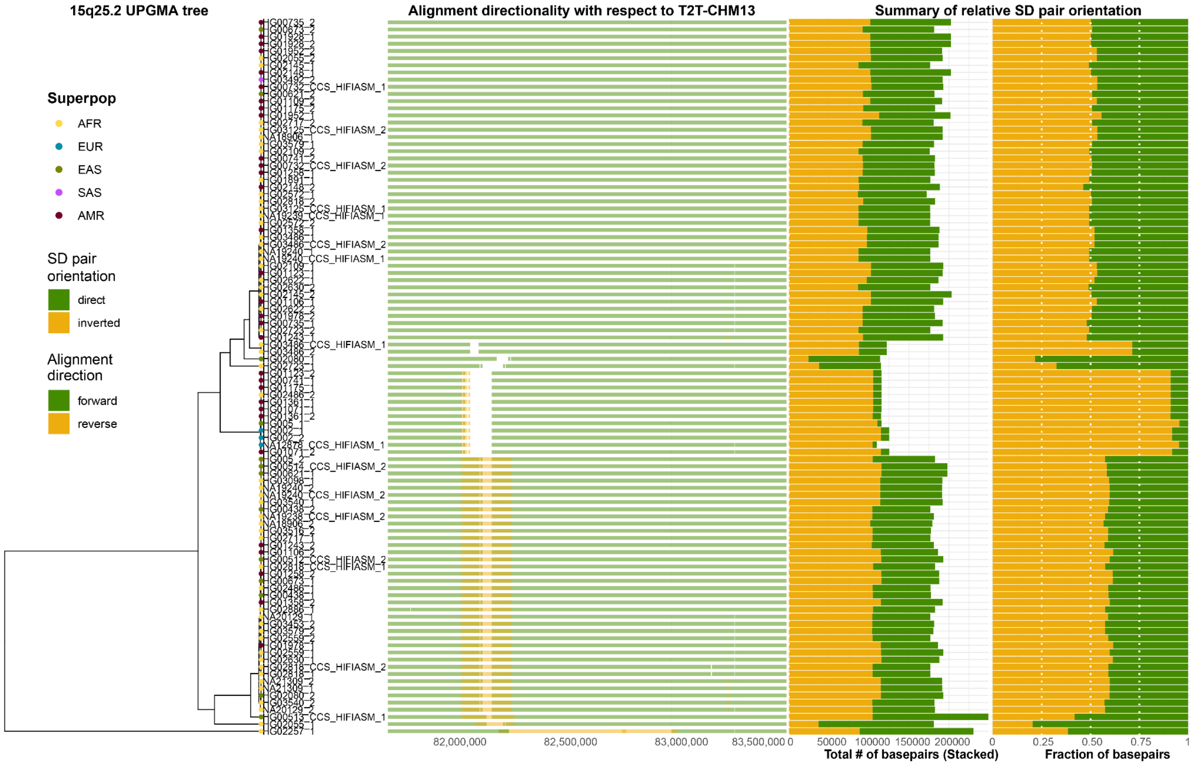


**Figure S16: Diverse structural haplotypes at 15q25.2 region.**
From left to right: (i) An UPGMA tree grouping complete assemblies (n=101) of 15q25.2 region into structurally similar groups based on their alignment to T2T-CHM13 (**Methods**). Superpopulation of origin for each sample is marked by colored dots. (ii) Visualization of alignment directionality (plus - green, minus - orange) of each assembly with respect to T2T-CHM13. (iii) Summary of the total number of base pairs for direct and reverse orientated SD pairs. (iv) Summary of the fraction of base pairs for direct and reverse orientated SD pairs.

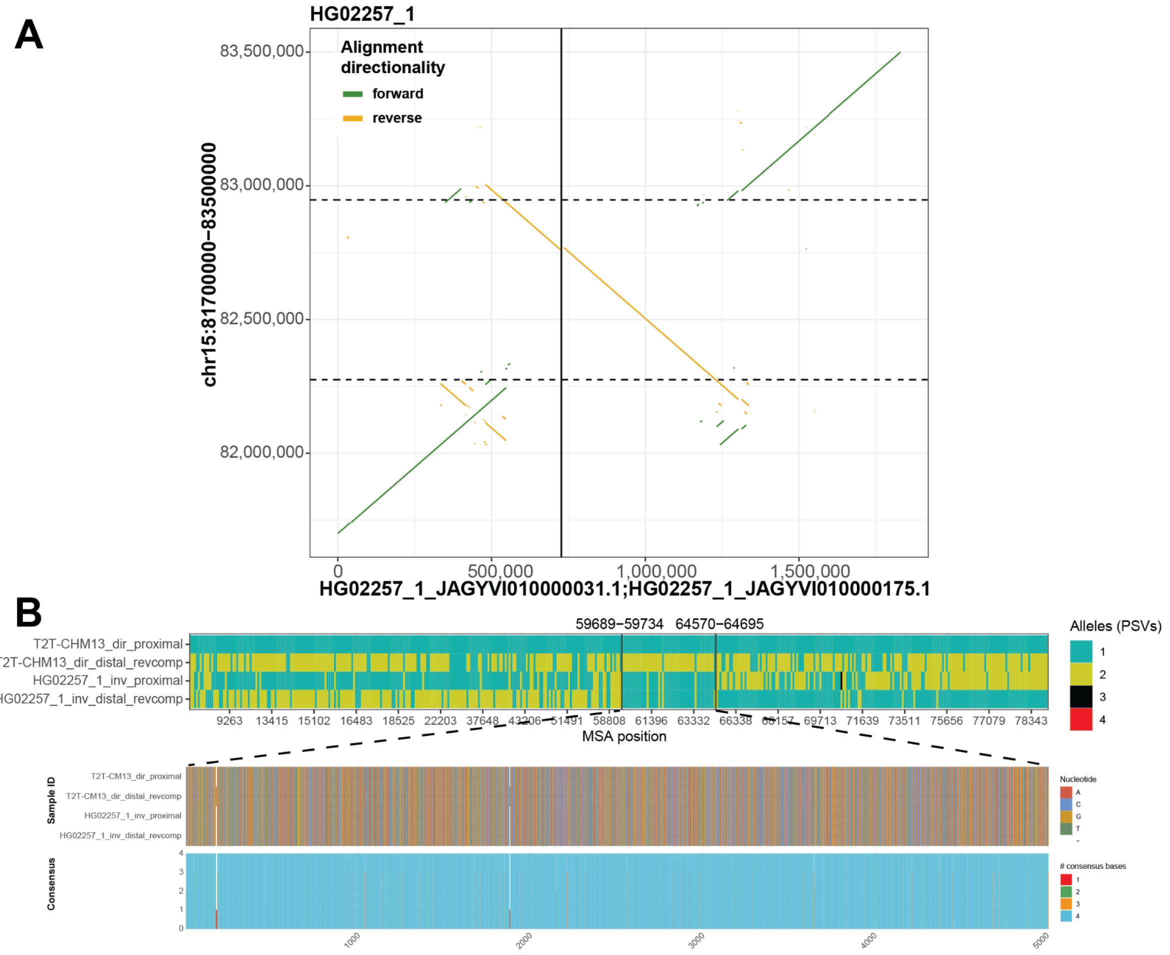


**Figure S17: Assembled inversion breakpoints at 15q25.2 and inversion breakpoint mapping.**
**A**) A dotplot showing the alignment directionality (yellow - reverse, green - forward) between HG02257 assembly of 15q25.2 region (x-axis) against T2T-CHM13 reference (y-axis). Reported inversion in T2T-CHM13 coordinates is highlighted by horizontal dashed lines. The position where one contig ends and another starts is marked by a solid vertical line. **B**) Visualization of multiple sequence alignment (MSA) between inversion flanking SDs from direct (T2T-CHM13) and inverted (HG02257) haplotype. Only paralog specific variants (PSVs) from the proximal (bright green) and distal (dark yellow) SDs are colored separately. Gaps in the MSA are colored white and alleles not present in the proximal and distal SDs are shown in black and red, respectively. Vertical solid lines depict detected change points, with numbers showing the change point position within flanking SDs. We predict that the inversion breakpoints lie between the 59,689 and 64,695 bp of flanking SDs. Below we zoom into ~5 kbp wide breakpoint region of high homology shown by almost perfect consensus across inversion flanking SDs.

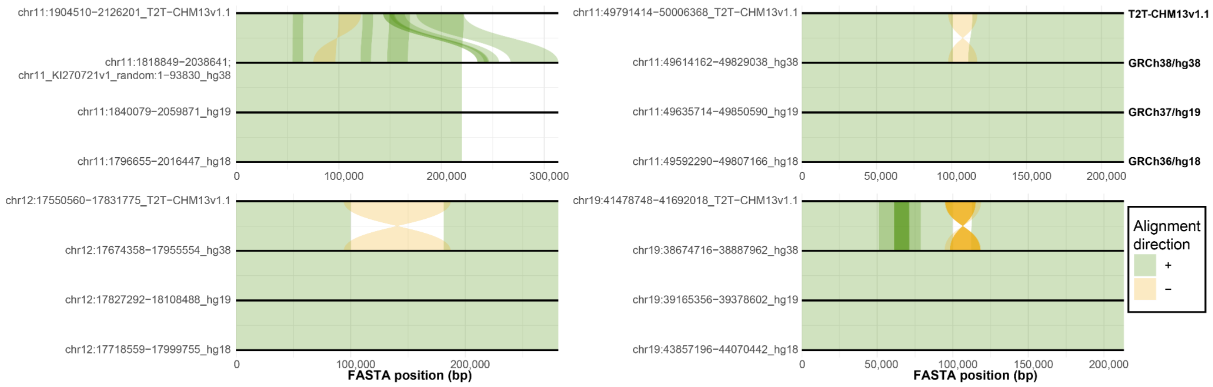


**Figure S18: Example of long-lasting misorientation errors in previous human genome references.**

Here we show a comparison of the FASTA sequences extracted from four misoriented regions (**Table S4**) across four versions of human genome reference assembly (from top to bottom: T2T-CHM13v1.1, GRCh38, GRCh37 and GRCh36). Alignment directionality is highlighted by direct ('+', green) and reverse ('-', orange) oriented flows between pairs of FASTA sequences.

#### SUPPLEMENTAL TABLES (S1-S5):

**Table S1: Nonredundant inversion callset reported in this study**
**Table S2: Putative novel inversions with respect to T2T-CHM13 reference**
**Table S3: Enrichment of inversion in pericentromeric regions**
**Table S4: List of minor alleles and resolved orientation errors in GRCh38**
**Table S5: Novel inversions in HPRC Strand-seq dataset**

#### Consortia:

**Human Genome Structural Variation Consortium (HGSVC):**
HGSVC co-chairs:
Charles Lee, Evan E. Eichler, Jan O. Korbel, Tobias Marschall

European Molecular Biology Laboratory (EMBL), Genome Biology Unit, Meyerhofstr. 1, 69117 Heidelberg, Germany
Jan O. Korbel, Bernardo Rodriguez-Martin, Tobias Rausch, Marc Jan Bonder, Wolfram Höps, Ashley D. Sanders, Benjamin Raeder, Patrick Hasenfeld, Oliver Stegle

Heinrich Heine University, Medical Faculty, Institute for Medical Biometry and Bioinformatics, Moorenstr. 20, 40225 Düsseldorf, Germany
Tobias Marschall, Peter Ebert, Jana Ebler, Hufsah Ashraf, Rebecca Serra Mari, Maryam Ghareghani

Department of Genome Sciences, University of Washington School of Medicine, Seattle, WA 98195, USA
Evan E. Eichler, David Porubsky, PingHsun Hsieh, Katherine M. Munson, William T. Harvey, Alexandra P. Lewis

The Jackson Laboratory for Genomic Medicine, Farmington, CT 06032, USA
Charles Lee, Christine Beck, Peter A. Audano, Qihui Zhu, Feyza Yilmaz, Pille Hallast

Department of Computational Medicine & Bioinformatics, University of Michigan, Ann Arbor, MI 48109, USA
Ryan E. Mills, Weichen Zhou

Institute for Genome Sciences, University of Maryland School of Medicine, Baltimore, MD 21201, USA
Scott E. Devine, Nelson T. Chuang, Luke J. Tallon

Center for Genomic Medicine, Massachusetts General Hospital, Department of Neurology, Harvard Medical School, Boston, MA 02114, USA
Michael E. Talkowski, Xuefang Zhao, Harrison Brand

Program in Computational Biology and Bioinformatics, Yale University, New Haven, CT 06520, USA
Mark B. Gerstein, Sushant Kumar

Xi’an Jiaotong University, Xi’an, Shaanxi, 710049, China
Kai Ye, Jiadong Lin, Xiaofei Yang

Department of Quantitative and Computational Biology, University of Southern California, Los Angeles, CA 90089, USA
Mark J.P. Chaisson, Jingwen Ren, Tsung-Yu Lu

Department of Computer & Information Sciences, Temple University, Philadelphia, PA 19122, USA
Xinghua Shi, Chong Li, Sky Gao, Bin Li, Chen Song

Department of Genetics, Yale School of Medicine, New Haven, CT 06510, USA
Ira M. Hall

Department of Genetics and Informatics Institute, School of Medicine, University of Alabama at Birmingham, Birmingham, AL 35294, USA
Zechen Chong, Yu Chen

European Molecular Biology Laboratory, European Bioinformatics Institute, Wellcome Genome Campus, Hinxton, Cambridge, CB10 1SD, United Kingdom
Sarah Hunt, Susan Fairley, Paul Flicek

New York Genome Center, New York, NY 10013, USA
Michael C. Zody, Wayne E. Clarke, Anna O. Basile, Marta Byrska-Bishop, André Corvelo, Uday S. Evani

Washington University, St. Louis, MO 63108, USA
Allison A. Regier, Haley J. Abel

University of Chicago, Chicago, IL 60637, USA
Yang I. Li, Zepeng Mu

Department of Bioinformatics and Computational Biology, The University of Texas MD Anderson Cancer Center, Houston, TX 77030, USA
Ken Chen

Genomes and Disease, Centre for Research in Molecular Medicine and Chronic Diseases (CIMUS), Universidade de Santiago de Compostela, Santiago de Compostela, Spain
Martin Santamarina, Jose M.C. Tubio

Bionano Genomics, San Diego, CA 92121, USA
Alex R. Hastie

Pacific Biosciences of California, Inc., Menlo Park, CA 94025, USA
Aaron M. Wenger

**Human Pangenome Reference Consortium (HPRC)**

Haley J. Abel[1],Lucinda L Antonacci-Fulton[2], Mobin Asri[3], Gunjan Baid[4], Carl A. Baker[5], Anastasiya Belyaeva[4], Konstantinos Billis[6], Guillaume Bourque[7,8,9], Silvia Buonaiuto[10], Andrew Carroll[4], Mark JP

Chaisson[11], Pi-Chuan Chang[4], Xian H. Chang[3], Haoyu Cheng[12,13], Justin Chu[12], Sarah Cody[2], Vincenza Colonna[10,14], Daniel E. Cook[4], Robert M. Cook-Deegan[15], Omar E. Cornejo[16], Mark Diekhans[3], Daniel Doerr[17], Peter Ebert[17], Jana Ebler[17], Evan E. Eichler[5,18], Jordan M. Eizenga[3], Susan Fairley[6], Olivier Fedrigo[19], Adam L. Felsenfeld[20], Xiaowen Feng[12,13], Christian Fischer[14], Paul Flicek[6], Giulio Formenti[19], Adam Frankish[6], Robert S. Fulton[2], Yan Gao[21], Shilpa Garg[22], Erik Garrison[14], Carlos Garcia Giron[6], Richard E. Green[23,24], Cristian Groza[25], Andrea Guarracino[26], Leanne Haggerty[6], Ira Hall[27,28], William T Harvey[5], Marina Haukness[3], David Haussler[3,18], Simon Heumos[29,30], Glenn Hickey[3], Kendra Hoekzema[5], Thibaut Hourlier[6], Kerstin Howe[31], Miten Jain[32], Erich D. Jarvis[33,18], Hanlee P. Ji[34], Alexey Kolesnikov[4], Jan O. Korbel[35], Jennifer Kordosky[5], Sergey Koren[36], HoJoon Lee[34], Alexandra P. Lewis[5], Heng Li[12,13], Wen-Wei Liao[2,37,27], Shuangjia Lu[27], Tsung-Yu Lu[38], Julian K. Lucas[3], Hugo Magalhães[17], Santiago Marco-Sola[39,40], Pierre Marijon[17], Charles Markello[3], Tobias Marschall[17], Fergal J. Martin[6], Ann McCartney[36], Jennifer McDaniel[41], Karen H. Miga[3], Matthew W. Mitchell[42], Jean Monlong[3], Jacquelyn Mountcastle[19], Katherine M. Munson[5], Moses Njagi Mwaniki[43], Maria Nattestad[4], Adam M. Novak[3], Sergey Nurk[36], Hugh E. Olsen[3], Nathan D. Olson[41], Benedict Paten[3], Trevor Pesout[3], Adam M. Phillippy[36], Alice B. Popejoy[44], David Porubsky[5], Pjotr Prins[14], Daniela Puiu[45], Mikko Rautiainen[36], Allison A Regier [2], Arang Rhie[36], Samuel Sacco[46], Ashley D. Sanders[47], Valerie A. Schneider[48], Baergen I. Schultz[20], Kishwar Shafin[4], Jonas A. Sibbesen[49], Jouni Sirén[3], Michael W. Smith[20], Heidi J. Sofia[20], Ahmad N. Abou Tayoun[50,51], Françoise Thibaud-Nissen[48], Chad Tomlinson[2], Francesca Floriana Tricomi[6], Flavia Villani[14], Mitchell R. Vollger[5,52], Justin Wagner[41], Brian Walenz[36], Ting Wang[53], Jonathan M. D. Wood[31], Aleksey V. Zimin[45,54], Justin M. Zook[41]

1 Division of Oncology, Department of Internal Medicine, Washington University School of Medicine, St. Louis, MO 63110, USA
2 McDonnell Genome Institute, Washington University School of Medicine, St. Louis, MO 63108, USA
3 UC Santa Cruz Genomics Institute, University of California, Santa Cruz, 1156 High St, Santa Cruz, CA, USA
4 Google LLC, 1600 Amphitheater Pkwy, Mountain View, CA 94043, USA
5 Department of Genome Sciences, University of Washington School of Medicine, Seattle, WA 98195, USA
6 European Molecular Biology Laboratory, European Bioinformatics Institute, Wellcome Genome Campus, Cambridge, CB10 1SD, UK
7 Department of Human Genetics, McGill University, Montreal, Québec H3A 0C7, Canada
8 Canadian Center for Computational Genomics, McGill University, Montreal, Québec H3A 0G1, Canada
9 Institute for the Advanced Study of Human Biology (WPI-ASHBi), Kyoto University, Kyoto 606-8501, Japan
10 Institute of Genetics and Biophysics, National Research Council, Naples 80111, Italy

11 University of Southern California, Quantitative and Computational Biology, Los Angeles, CA, USA
12 Department of Data Sciences, Dana-Farber Cancer Institute, Boston, MA 02215, USA
13 Department of Biomedical Informatics, Harvard Medical School, Boston, MA 02215, USA
14 Department of Genetics, Genomics and Informatics, University of Tennessee Health Science Center, Memphis, TN 38163, USA
15 Arizona State University, Barrett & O'Connor Washington Center, Washington DC, USA
16 School of Biological Sciences, Washington State University, Pullman WA 99163, USA
17 Institute for Medical Biometry and Bioinformatics, Medical Faculty, Heinrich Heine University Düsseldorf, Düsseldorf, Germany
18 Howard Hughes Medical Institute, Chevy Chase, MD 20815, USA
19 The Vertebrate Genome Laboratory, The Rockefeller University, New York, NY 10065, USA
20 National Institutes of Health (NIH)–National Human Genome Research Institute, Bethesda, MD, USA
21 Center for Computational and Genomic Medicine, The Children's Hospital of Philadelphia, Philadelphia, PA 19104, USA.
22 Department of Biology, University of Copenhagen, Denmark
23 Department of Biomolecular Engineering, University of California, Santa Cruz, 1156 High St., Santa Cruz, CA 95064, USA
24 Dovetail Genomics, Scotts Valley, CA 95066, USA
25 Quantitative Life Sciences, McGill University, Montreal, Québec H3A 0C7, Canada
26 Genomics Research Centre, Human Technopole, Milan 20157, Italy
27 Department of Genetics, Yale University School of Medicine, New Haven, CT 06510, USA
28 Center for Genomic Health, Yale University School of Medicine, New Haven, CT 06510, USA
29 Quantitative Biology Center (QBiC), University of Tübingen, Tübingen 72076, Germany
30 Biomedical Data Science, Department of Computer Science, University of Tübingen, Tübingen 72076, Germany
31 Tree of Life, Wellcome Sanger Institute, Hinxton, Cambridge, CB10 1SA, UK
32 Northeastern University, Boston, MA 02115, USA
33 The Rockefeller University, New York, NY 10065, USA
34 Division of Oncology, Department of Medicine, Stanford University School of Medicine, Stanford, CA, 94305, USA
35 European Molecular Biology Laboratory, Genome Biology Unit, Meyerhofstr. 1, 69117 Heidelberg, Germany
36 Genome Informatics Section, Computational and Statistical Genomics Branch, National Human Genome Research Institute, National Institutes of Health, Bethesda, MD 20892, USA
37 Department of Medicine, Washington University School of Medicine, St. Louis, MO 63110, USA
38 University of Southern California, Quantitative and Computational Biology, 3551 Trousdale, Pkwy, Los Angeles, CA, USA
39 Computer Sciences Department, Barcelona Supercomputing Center, Barcelona, Spain

40 Departament d'Arquitectura de Computadors i Sistemes Operatius, Universitat Autònoma de Barcelona, Barcelona, Spain
41 Material Measurement Laboratory, National Institute of Standards and Technology, Gaithersburg, MD 20877, USA
42 Coriell Institute for Medical Research, Camden, NJ 08103, USA
43 Department of Computer Science, University of Pisa, Pisa 56127, Italy
44 Department of Public Health Sciences, University of California, Davis, One Shields Avenue, Medical Sciences 1C, Davis, CA 95616
45 Department of Biomedical Engineering, Johns Hopkins University, Baltimore 21218, MD, USA
46 Department of Ecology & Evolutionary Biology, University of California, Santa Cruz, 1156 High St, Santa Cruz, CA, USA
47 Berlin Institute for Medical Systems Biology, Max Delbrück Center for Molecular Medicine in the Helmholtz Association, Berlin, Germany
48 National Center for Biotechnology Information, National Library of Medicine, National Institutes of Health, Bethesda, MD 20894, USA
49 Center for Health Data Science, University of Copenhagen, Denmark
50 Al Jalila Genomics Center of Excellence, Al Jalila Children's Specialty Hospital, Dubai, UAE
51 Center for Genomic Discovery, Mohammed Bin Rashid University of Medicine and Health Sciences, Dubai, UAE
52 Division of Medical Genetics, University of Washington School of Medicine, Seattle, WA 98195, USA
53 Department of Genetics, Washington University School of Medicine, St. Louis, MO 63110, USA
54 Center for Computational Biology, Johns Hopkins University, Baltimore, MD 21218, USA
